# Physiological Roles of Carbonic Anhydrases and an MpsAB-like Bicarbonate Transporter in *Bacillus subtilis*

**DOI:** 10.64898/2026.09.08.750065

**Authors:** Grayson Barnes, John D. Helmann

**Author notes:** Address correspondence to: John D. Helmann.

## Abstract

Carbonic anhydrase (E.C. 4.2.1.1) is an enzyme that catalyzes the reversible hydration of CO_2_ to carbonic acid (H_2_CO_3_), which dissociates to bicarbonate (HCO_3_^−^) at intracellular pH. Copious amounts of CO_2_ are produced by catabolism, although much of this is lost by diffusion from the cell. In *Bacillus subtilis*, anabolic processes rely largely on bicarbonate as a substrate rather than CO_2_. While CO_2_ reacts spontaneously with water to yield bicarbonate, the rate of this reaction is too slow to keep up with cell requirements. *B. subtilis* encodes three putative β-class carbonic anhydrases, here renamed *canA*(*yvdA)*, *canB*(*ytiB*), and *canC*(*ybcF)*. The *canC* gene is encoded in an operon with *ndhF*-*mpsB*(*ybcC)*, which encodes a candidate MpsAB-type bicarbonate transporter. Here we demonstrate that a strain lacking *canA, canB, canC,* and *mpsB* (Δ4) has a severe growth defect at atmospheric CO_2_. This defect can be overcome by growing cells with supplemental CO_2_ or by plating at high cell density. We isolated suppressors of Δ4 and identified mutations in *resD* that suppress the requirement for supplemental CO_2_ to support growth. ResD functions as a global regulator of genes important for both aerobic and anaerobic respiration. We demonstrate that a *resD* null mutation results in metabolic changes that lead to an increased generation of CO_2_. We infer that this results in an increase in spontaneous bicarbonate formation that is sufficient to support cell growth. We conclude that the requirement for bicarbonate concentrating mechanisms may be bypassed under conditions that increase endogenous CO_2_ generation.

**Importance:** Carbonic anhydrase (CA) catalyzes the reversible hydration of CO_2_ to bicarbonate. This enzyme is found in all domains of life and has evolved multiple times. In heterotrophic organisms, CA is important for concentrating bicarbonate for anabolic reactions such as fatty acid, amino acid, and menaquinone synthesis. Bicarbonate can also be directly imported by dedicated transporters. Free-living bacteria are often unable to grow in the absence of a bicarbonate concentrating mechanism, and CAs and bicarbonate transporters are also important in pathogenesis. Thus, understanding how cells obtain bicarbonate may lead to the development of novel antibiotics and highlight new therapeutic targets.

## Introduction

Carbonic anhydrase (CA) is a metalloenzyme that catalyzes the reversible hydration of dissolved CO_2_ to carbonic acid. This activity was first described in 1933 by Meldrum and Roughton after an observation by Henriques that the presence of blood greatly accelerated the hydration of CO_2_ (1). CA is ubiquitous in mammalian tissues, plants, and unicellular green algae, with a more sporadic distribution noted in the Bacterial and Archaeal domains (2). CA has evolved multiple times with eight structurally distinct families identified to date (α, β, γ, δ, ζ, η, θ, ι). A recent phylogenomic survey of the three dominant CA classes (α, β, and γ) identified representatives in ∼70% of Bacteria and 83% of Archaea (3). Within cultured bacterial representatives (∼15,000), 86% harbor at least one CA. Remarkably, most organisms encode multiple CA isozymes ranging from several (1.8 ±0.9 in Archaea and 2.8 ± 1.6 in Bacteria) to many (10 or more in Fungi and Algae) (3). This ubiquity attests to the physiological importance of CA in diverse organisms.

CA was first appreciated for its role in mammalian respiration where it hydrates CO_2_ released from the tissues to bicarbonate to allow for efficient transport through the bloodstream to the lungs. At the lungs, the reverse reaction facilitates the rapid generation of CO_2_ for exhalation (4). CA is also widely heralded for its role in photosynthesis. Carbon fixation via the Calvin cycle begins with ribulose-1,5-bisphosphate carboxylase/oxygenase (RuBisCo) which requires elevated concentrations of CO_2_ to efficiently fix carbon into organic matter (5). CA and bicarbonate transporters are key players in the carbon-concentrating mechanisms that help supply RuBisCo with high concentrations of CO_2_.

The role of CA and CA homologs is less studied in non-photosynthetic Bacteria. *Escherichia coli* has two β-class CA isozymes, Can and CynT. Can is essential for growth at atmospheric levels of CO_2_, and CA function can also be provided by the cyanate-inducible CynT enzyme (6). Insights into the essential role of Can at atmospheric CO_2_ levels were obtained by considering: 1) rates of production of CO_2_ from catabolism, 2) loss of CO_2_ by diffusion, and 3) the amount of bicarbonate required to support anabolism (6). Since CO_2_ can rapidly diffuse across the membrane, hydration to bicarbonate serves to trap inorganic carbon within the cell and to supply bicarbonate to anabolic enzymes. In the absence of CA, it was calculated that the intracellular concentration of CO_2_ is ∼10^3^-fold too low to meet cell demands (6).

Consideration of those bacteria that do not encode CA is also instructive. For example, *Staphylococcus aureus* lacks CA and instead relies on an energy-dependent bicarbonate transporter (MpsAB) (7, 8). Phylogenomic comparisons reveal that many bacteria within the Bacillota contain MpsAB instead of, or in addition to, CA (8). A lack of CA is also seen in bacteria that require high levels of CO_2_ for growth (capnophilic organisms). These include some syntrophic bacteria that reside in communities, as seen in *Symbiobacterium thermophilum* (9), and many that are host associated. In the case of *Brucella* spp. a requirement for elevated CO_2_ to support growth is correlated with the presence of a CA pseudogene (10–12).

*Bacillus subtilis* encodes three putative β-class CAs, but their functions have not been previously investigated. We here set out to establish the physiological importance of these three CA genes, which we have renamed as *canA*(*yvdA)*, *canB*(*ytiB*), and *canC*(*ybcF)*. We demonstrate that these three genes are individually and collectively dispensable for growth at atmospheric levels (∼0.043%) of CO_2_. This phenotype revealed the role of a predicted MpsAB-type bicarbonate transporter in *B. subtilis*. A quadruple mutant strain (Δ4) that lacks all three CA genes (*canA, canB* and *canC*) and *mpsB* (formerly *ybcC*) is unable to grow as single colonies without CO_2_ supplementation. We surmise that the flux of CO_2_ from catabolism and the slow rate of spontaneous generation of bicarbonate is insufficient to support the key bicarbonate-dependent anabolic reactions that support cell growth at low density. Consistent with this notion, we recovered Δ4 suppressor strains carrying mutations in *resD* that recover growth without CO_2_ supplementation. ResD is a two-component response regulator that controls the transcription of key genes for respiration. We establish that this suppressor strain (Δ4 Δ*resD*) had a higher rate of CO_2_ production, which was sufficient to allow growth of single colonies at atmospheric levels of CO_2_. Although suppressors have been sporadically reported that improve the growth of CA-deficient cells, our results provide molecular insights into the types of genetic changes that may be responsible.

## Results

### *Bacillus* subtilis has redundant bicarbonate concentrating systems

*B. subtilis* 168 has three genes that code for proteins with sequence and predicted structural similarity to β-class CA enzymes: *canA* (formerly *yvdA*), *canB* (formerly *ytiB*), and *canC* (formerly *ybcF*) (Fig. 1). We hypothesized that at least one of these putative CA enzymes would be required to support cell growth under atmospheric levels (∼0.043%) of CO_2_. However, we were easily able to construct a *canA canB canC* (ΔABC) triple mutant under atmospheric CO_2_ conditions. We note that the atmospheric CO_2_ concentration in these studies was that of a well-ventilated research laboratory, which on average was ∼10% higher than outdoor air (∼472+39 ppm; n=20; Table S1). The construction of multiple mutant strains with strong synthetic growth defects by chromosomal DNA transformation often results in congression. When congression occurs, the recovered transformants acquire the selectable marker associated with the new gene deletion as well as a WT copy of one of the genes that has already been deleted. However, very little congression was observed during construction of the ΔABC strain, and all three gene deletions were confirmed by PCR. This led us to suspect that there may be a fourth unannotated CA in the *B. subtilis* genome.

**Figure 1.**
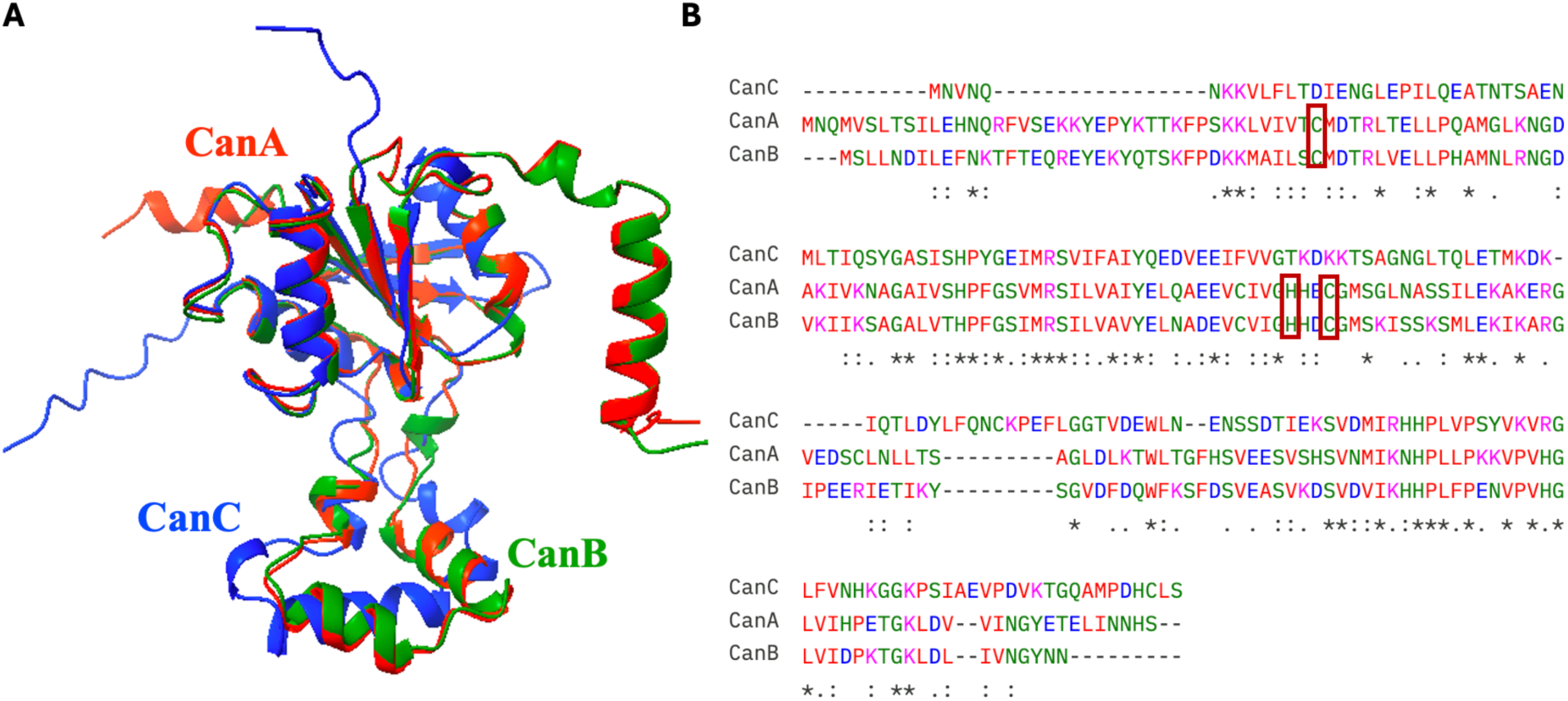
*Bacillus subtilis* has three predicted carbonic anhydrases. a) predicted structures of *B. subtilis* carbonic anhydrases CanA (red), CanB (green) and CanC (blue) aligned using matchmaker command in ChimeraX. b) Multiple sequence alignment of *B. subtilis* CanA, CanB, and CanC aligned with MUSCLE (EMBL server). A red box is placed around the predicted Zn^2+^ binding residues in CanA and CanB.

To search for a possible fourth β-class CA we used HHPred to find more distantly related proteins. In addition to the three CAs, we found another protein with very weak similarity to CAs (Fig. S1). YbcC is a homolog of MspB (46% identity), a subunit of the bicarbonate transporter found in *Staphylococcus aureus* (MpsAB) (8) (Fig. S2). Consistent with this assignment, *ybcC* is directly downstream of *ndhF*, which codes for a protein that shares sequence similarity with the MpsA subunit of the *S. aureus* bicarbonate transporter. In *S. aureus*, deletion of *mpsB* leads to a severe growth defect that can be overcome by complementing back the bicarbonate transporter, by expressing an *E. coli* carbonic anhydrase, or by supplementation with CO_2_ (7, 8).

To test if this putative bicarbonate transporter (*mpsAB*) was supporting growth of the ΔABC strain at atmospheric levels of CO_2_, we transformed a *mpsBcanC*::*erm* mutation into the Δ*canA* Δ*canB* background to generate a Δ*canA* Δ*canB mpsBcanC*::*erm* (Δ4) mutant strain. This mutant was difficult to construct, with frequent congression that led to replacement of one of the pre-existing CA deletions with a WT copy of the gene. This suggests that there is a high selective pressure to keep at least one carbonic anhydrase or a bicarbonate transporter in the genome. We did recover one Δ4 clone that could be re-streaked and grew, albeit poorly, in regions of high initial cell density, but not as single colonies (Fig. S3). The strain was validated by whole-genome sequencing, which identified all four expected deletion mutations and no additional second-site suppressor mutations when compared with the reference 168 WT genome. When plated on LB agar to recover single colonies, the Δ4 strain was only able to grow in the presence of elevated levels of CO_2_. When incubated in a 1.2 L air-tight plastic container small colonies were observed at ∼0.5% CO_2_ and much larger colonies with 5% CO_2_ (Fig. 2). We conclude that *B. subtilis* requires at least one CA or a bicarbonate transporter for growth with atmospheric levels of CO_2_.

**Figure 2.**
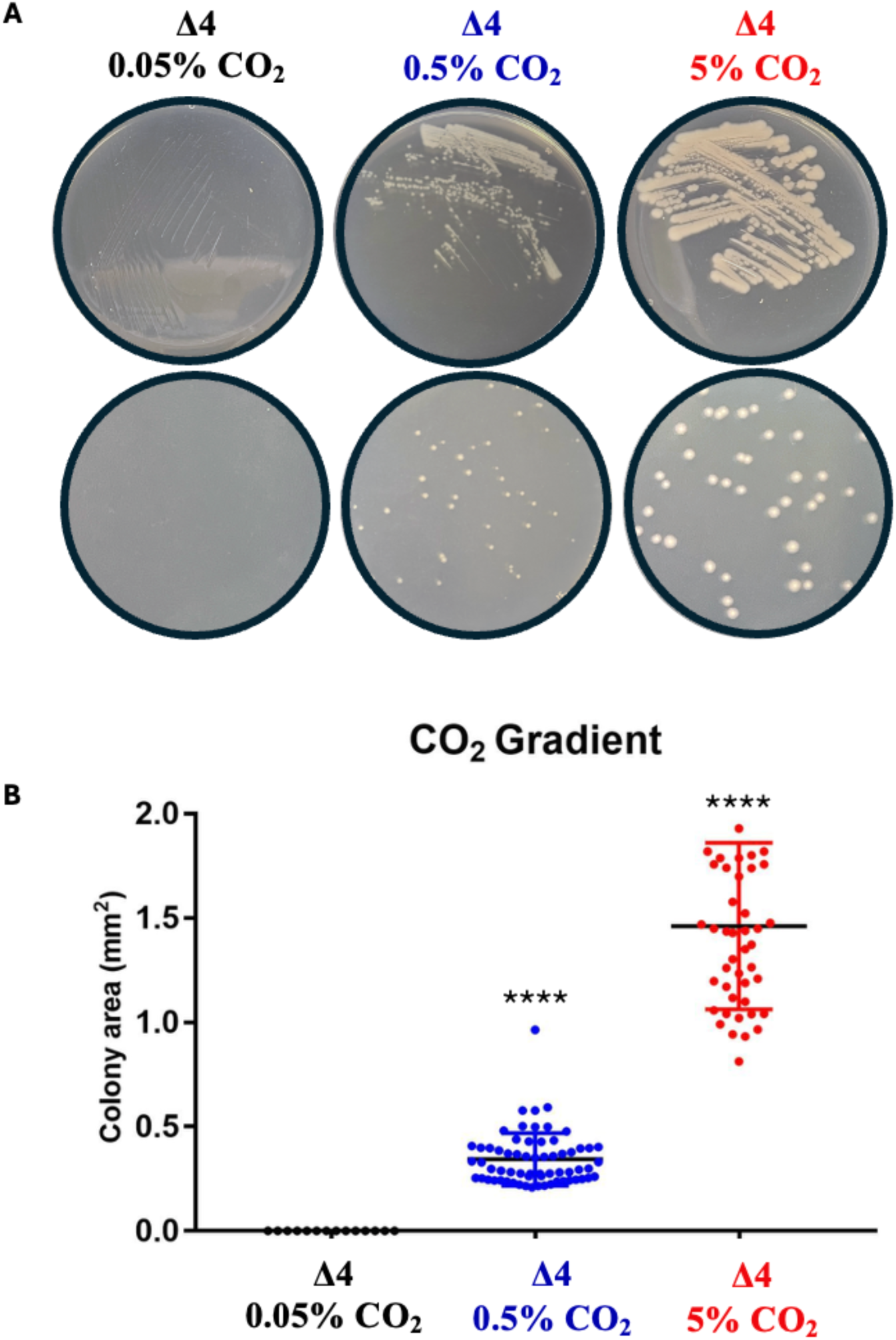
Growth of the Δ4 strain with and without supplemental CO_2_. a) Streaks (top) and colony size (bottom) of the Δ4 strain grown at 0.05%, 0.5%, and 5% CO_2_. b) Quantification of colony size of Δ4 grown at 0.05%, 0.5%, and 5% CO_2_. P-values were calculated using one-way ANOVA with Tukey’s multiple comparisons, **** p<0.0001.

### CanB is the major CA isozyme on LB agar medium

By comparison of the Δ4 and Δ3 strains, we infer that the MpsAB bicarbonate importer is sufficient to support growth even in the absence of all three genes encoding β-CA homologs. Next, we wished to test which of the three putative CA enzymes could individually support growth. We therefore generated all possible triple mutant strains lacking MpsB and two of the three CA homologs. Like the Δ4 strain, the *canB canC mpsB* triple mutant was unable to form colonies at atmospheric levels of CO_2_ (Figs. 3A, 3B). However, slow growth was observed when cells from a frozen glycerol stock were streaked at high density (Fig. S3). We conclude that CanA is insufficient to support growth of single colonies under these conditions. In contrast, CanB and CanC were individually sufficient to support growth without supplemental CO_2_ (Fig. 3C). As expected, growth of all mutant strains was improved when grown under a 5% CO_2_ atmosphere (Figs. 3B, 3D).

**Figure 3.**
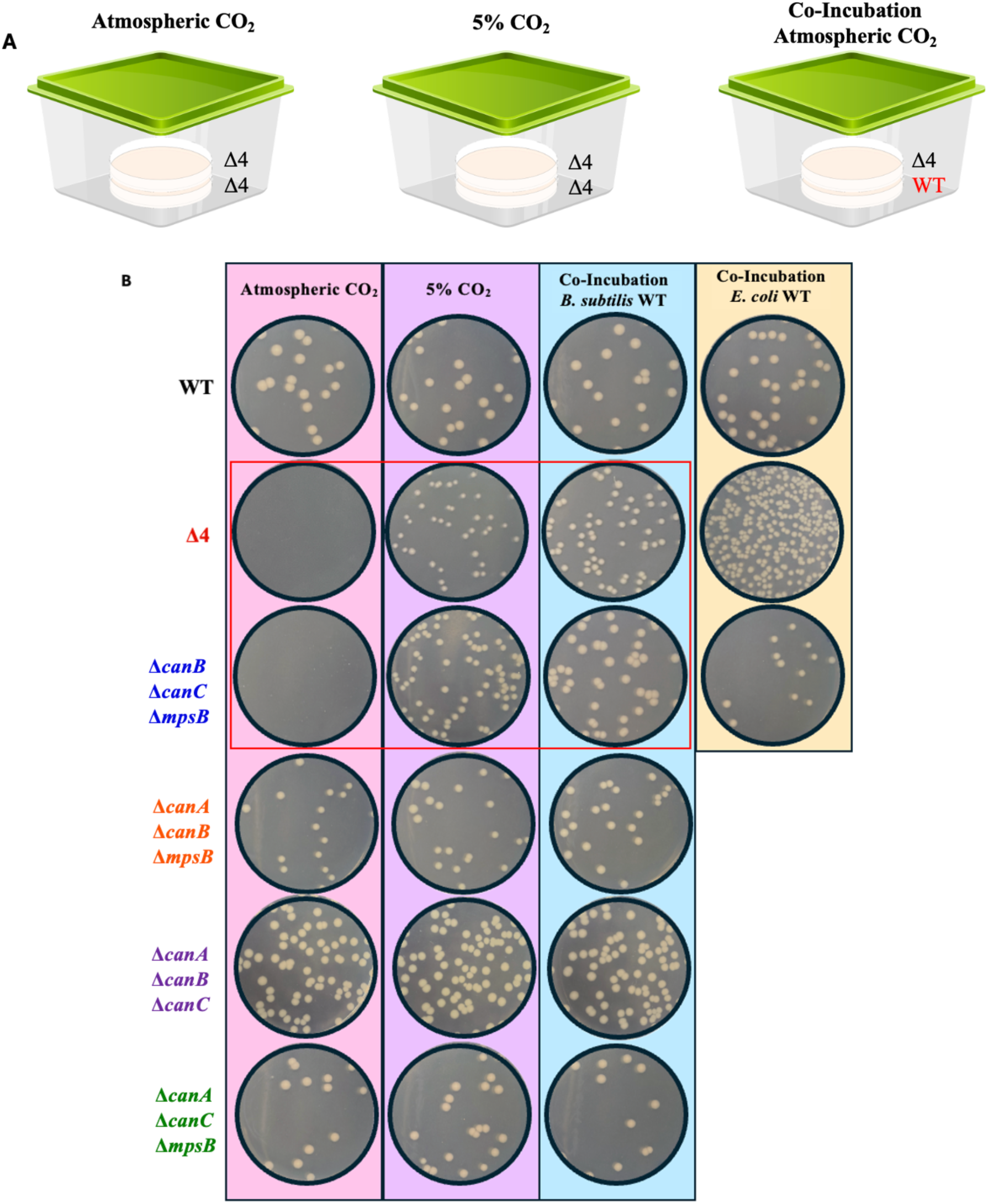
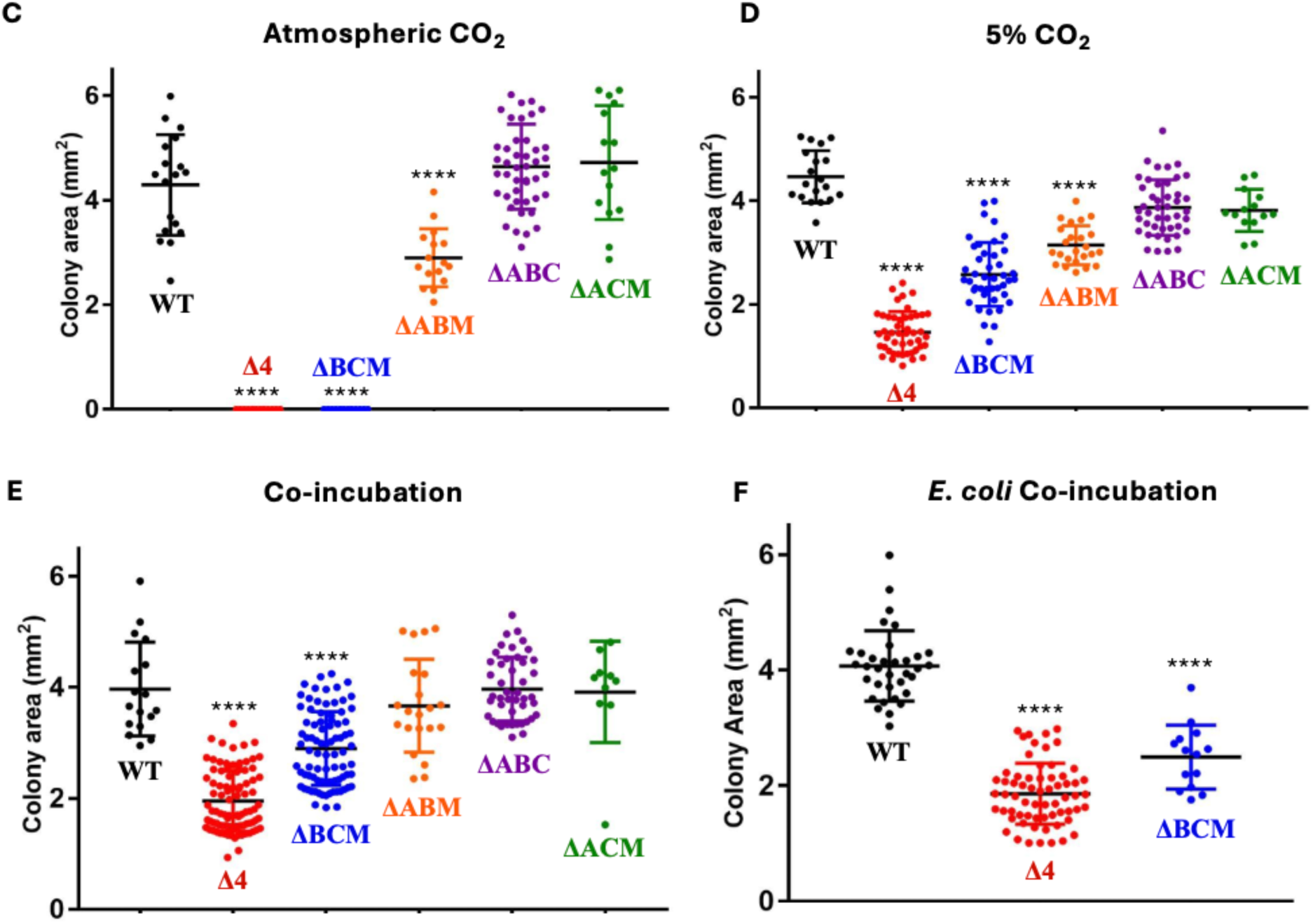
Growth of Δ4 and Δ*canB* Δ*canC* Δ*mpsB* can be rescued by elevated CO_2_. a) Representation of experimental set up. ∼1.2 L airtight plastic containers were used to grow the strains under a define starting atmosphere. b) Well-isolated colonies of WT, Δ4, Δ*canB* Δ*mpsBcanC* (BCM), Δ*canA* Δ*canB* Δ*mpsB* (ABM), Δ*canA* Δ*canB* Δ*canC* (ABC), Δ*canA* Δ*mpsBcanC* (ACM) were analyzed for size using ImageJ software. Strains were grown on LB agar plates grown at 37° C at either room atmosphere (atmospheric CO_2_ ∼0.047%), 5% CO_2_, or co-incubated with a lawn of WT *B. subtilis* or *E. coli*. Plates were imaged after 19 hours. Figure is representative of 3 independent biological replicates. c) Quantification of colony size of WT, Δ4, Δ*canB* Δ*mpsBcanC* (BCM), Δ*canA* Δ*canB* Δ*mpsB* (ABM), Δ*canA* Δ*canB* Δ*canC* (ABC), Δ*canA* Δ*mpsBcanC* (ACM) at c) atmospheric CO_2_, d) 5% CO_2_, and e) co-incubated with *B. subtilis*. f) Quantification of colony size of WT, Δ4 and Δ*canB* Δ*mpsBcanC* co-incubated with *E. coli*. Note some values have been repeated for clarity. P-values were calculated using one-way ANOVA with Tukey’s multiple comparisons test. Each mutant was compared WT grown in that same condition, **** p<0.0001

We extended these studies by constructing and testing single and double mutant strains (Figs. 4, S3). As expected from our initial work, all strains with single gene deletions were able to form single colonies on LB agar, with the *mpsB* mutant having a small but statistically significant growth defect. Interestingly, the *canB* and *canC* mutants appeared to form somewhat larger colonies than WT (Fig. 4B). We have not further investigated these modest growth effects. More striking, however, was the severe growth defect noted for the *canB mpsB* double mutant (Fig. 4). While a mild growth defect was observed when grown densely in a streak, this strain was unable to form single colonies and could only be partially rescued by elevated levels of CO_2_ (Fig. 4 and S3). This suggests that the two remaining CA-encoding genes (*canA* and *canC*) are insufficient to support growth as single colonies. Taken together, these growth observations support the hypothesis that CanB is the major CA under these conditions, with CanC playing a secondary role. We note, however, that the severe growth defect of the *canB mpsB* double mutant (Fig. 4) conflicts with the relatively modest growth defect of the triple mutant additionally lacking CanA (*canA canB mpsB*; Fig. 3). One possible explanation is a compensatory up-regulation of *canC* in the triple mutant strain. Indeed, we noted an ∼2-fold increase in *canC* mRNA levels in the *canA canB mpsB* triple mutant compared to WT (Fig. S4).

**Figure 4.**
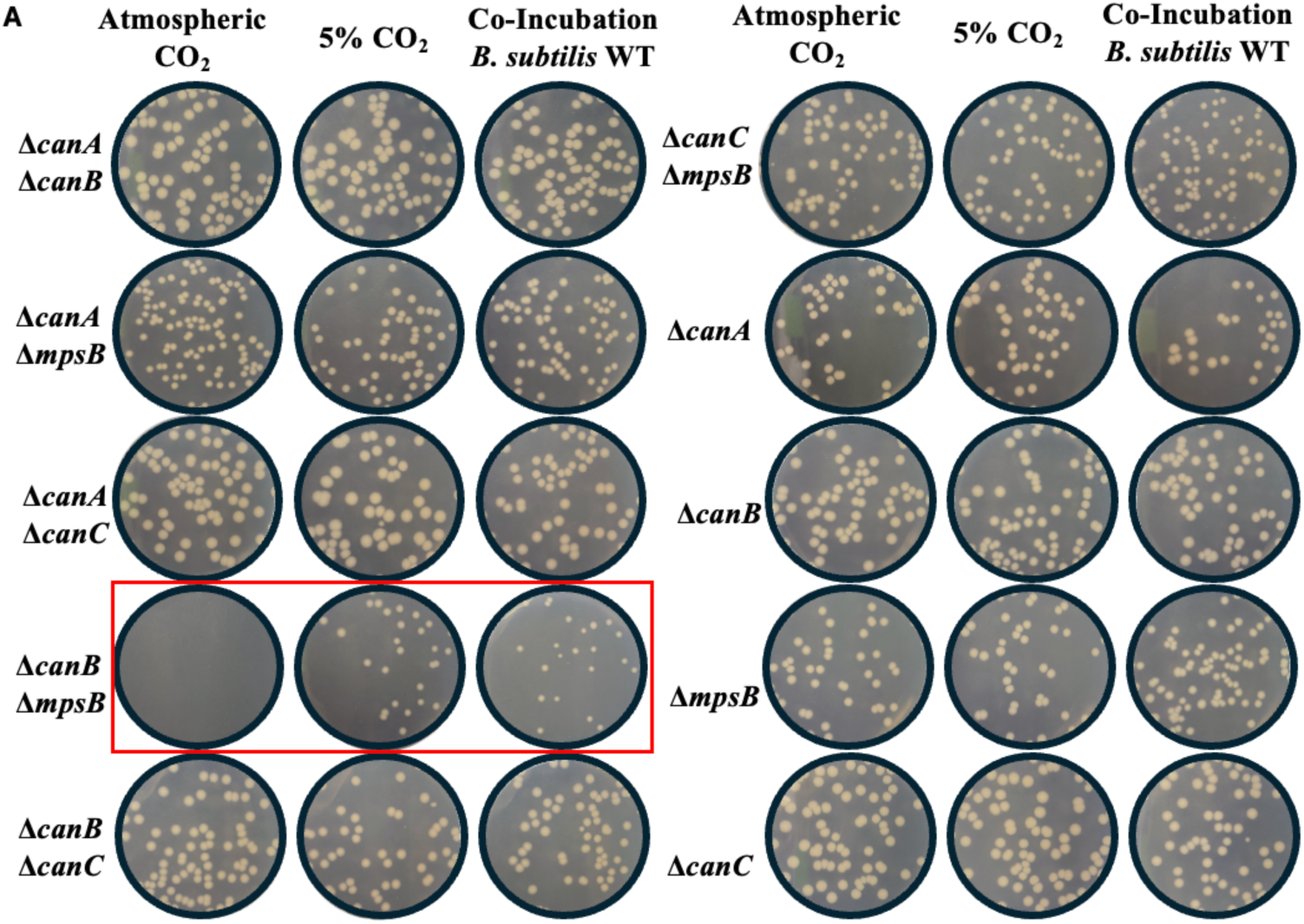
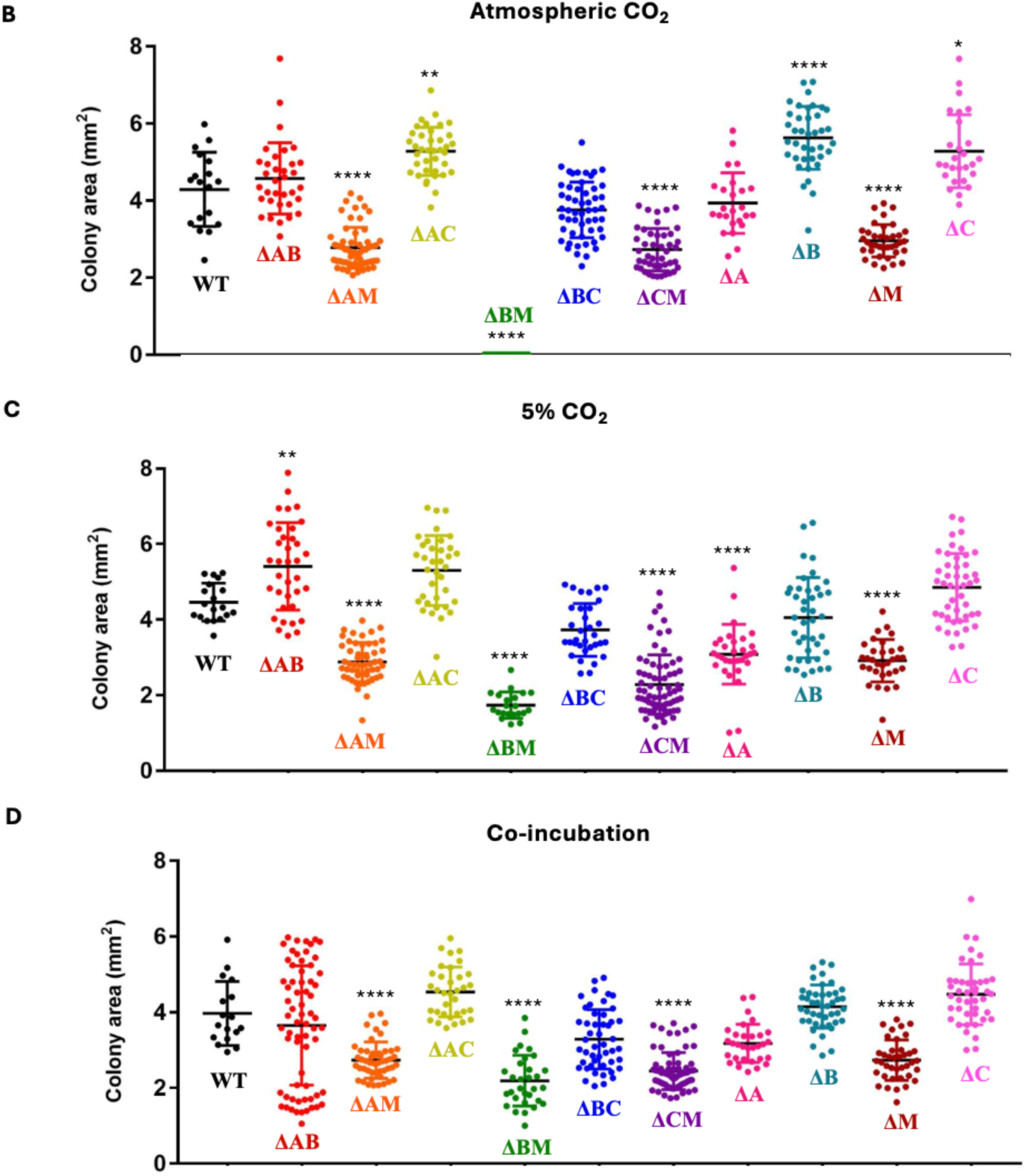
Colony size of Δ*canB* Δ*mpsB* can be rescued by elevated CO_2_. a) Well isolated colonies of WT, Δ*canA* Δ*canB* (ΔAB), Δ*canA* Δ*mpsB* (ΔAM), Δ*canA* Δ*canC* (ΔAC), Δ*canB* Δ*mpsB* (ΔBM), Δ*canB* Δ*canC* (ΔBC), Δ*mpsBcanC* (ΔCM), Δ*canA* (ΔA), Δ*canB* (ΔB), Δ*mpsB* (ΔM), and Δ*canC* (ΔC) were analyzed for size using ImageJ software. Strains were grown on LB agar plates grown at 37°C at CO_2_, 5% CO_2_, and co-incubated with a lawn of WT *B. subtilis*. Plates were imaged after 19 hours. Figure is representative of 3 independent biological replicates. Quantification of colony size of WT, Δ*canA* Δ*canB*, Δ*canA* Δ*mpsB*, Δ*canA* Δ*canC*, Δ*canB* Δ*mpsB*, Δc*anB* Δ*canC*, Δ*mpsBcanC*, Δ*canA*, Δ*canB*, Δ*mpsB*, and Δ*canC* at b) atmospheric CO_2_, c) 5% CO_2_, and d) co-incubated with a lawn of WT *B. subtilis* P-values were calculated using one-way ANOVA with Tukey’s multiple comparisons test. Each mutant was compared WT grown in that same condition, **** p<0.0001, ** p<0.01, and * p<0.05.

### Growth defects in bicarbonate deficient strains can be restored through co-incubation

If the growth defect of mutant strains lacking the bicarbonate importer and one or more CA enzymes is due to a deficiency in CO_2_/bicarbonate, we would expect that CO_2_ released from other respiring bacteria might rescue growth. Using the same ∼1.2 L air-tight plastic container system used for colony size measurements, we tested the ability of a growing lawn of either *B. subtilis* or *E. coli* to rescue growth of the compromised strains (Fig. 3, 4). Like cells incubated at 5% CO_2_, incubation with a growing lawn of cells efficiently rescued growth of cells in containers at atmospheric CO_2_ (generally <0.05%). To determine the amount of CO_2_ generated under these growth conditions, we measured the net release of CO_2_ in containers with a growing lawn of WT*B. subtilis*. Under our growth conditions, we recovered 70 mg (wet weight) of bacterial cells from lawns grown for 19 h at 37 °C. The final CO_2_ concentration in the container after growth was increased 100-fold to 4.7±0.3% (n=3), which corresponds to a net release of ∼100 mg CO_2_ (Table S1).

### Chemical complementation fails to rescue the growth of bicarbonate deficient mutants

Previous studies suggest that the inability of CA-deficient heterotrophs to grow with atmospheric levels of CO_2_ is likely due to a failure in one or more critical, bicarbonate-dependent enzymes (6). These include key enzymes necessary for fatty acid synthesis (acetyl-CoA carboxylase), amino acid synthesis (carbamoyl-phosphate synthase; 5-aminoimidazole ribotide carboxylase), the anaplerotic enzyme pyruvate carboxylase, and biotin carboxylase. Bicarbonate also functions in menaquinone biosynthesis by serving as a catalytic cofactor for 1,4-dihydroxy-2-naphthoyl-CoA synthase (MenB) (13). In *Saccharomyces cerevisiae*, a CA-deficient strain could be complemented for growth by supplementation with fatty acids, uracil, aspartate, and arginine (14). However, it has proven difficult to chemically complement CA-deficient mutants in heterotrophic bacteria (6).

To determine if we could bypass the requirement for CA or bicarbonate import by supplementation we amended our growth medium with uracil, adenine, proline, aspartate, the menaquinone precursor 1,4-dihydroxy-2-naphthoic acid (DHNA), and the fatty acids palmitic acid and 12-methyltetradecanoic acid (Fig. S5). In *E. coli*, fatty acid synthesis imposes a large requirement for bicarbonate (6). In *B. subtilis*, supplementation with palmitic acid and 12-methyltetradecanoic acid has been reported to suppress the inhibition of fatty acid synthesis by cerulenin (15), although fatty acid degradation limits incorporation (16). In our studies, amendment with these nutrients, alone and in combination, was unable to restore the growth of the Δ4 strain (Fig. S5). This negative result suggests that the concentrations used may not have been optimal, *Bacillus* may not efficiently uptake these nutrients and incorporate them into the cell, or perhaps there are other unrecognized bicarbonate-dependent processes.

### Expression of bicarbonate concentrating systems in different growth media

The presence of redundant systems to support bicarbonate-dependent metabolism may be due, in part, to differences in their expression or regulation. Prior work, using tiling arrays to monitor gene expression across the *B. subtilis* genome as a function of growth conditions (16), reveals that *canB* is expressed about 10-fold higher than *canA*, *canC*, and *mpsB* across most surveyed growth conditions (Fig. 5A). This is consistent with our inference that CanB is the major CA enzyme on LB agar plates. However, there were notable exceptions. Intriguingly, *canA* is most highly expressed when cells were grown as confluent lawns on LB medium. In addition, the *mpsAB* importer and co-transcribed *canC* are highly expressed under heat shock conditions. The details of these measurements can be interrogated using the *B. subtilis* expression database browser through the SubtiWiki website (17).

**Figure 5.**
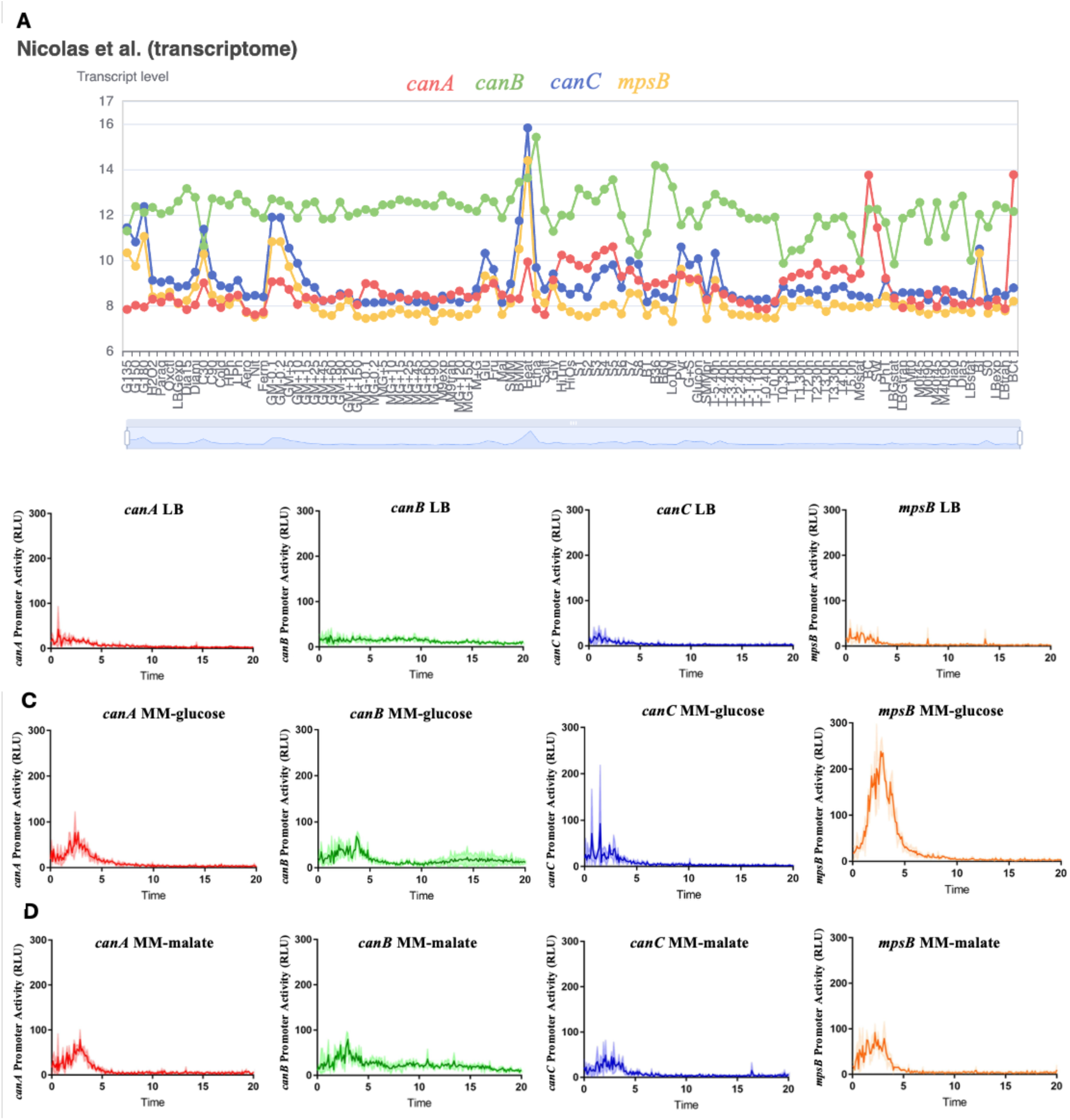
*mpsB* expression spikes during growth in MM-glucose. a) Expression of *canA*, *canB*, *canC*, and *mpsB* in many media and stress conditions as reported in the expression database browser on SubtiWiki. Expression of *canA*, *canB*, *canC*, and *mpsB* in b) LB, c) MM-glucose, and d) MM-malate monitored using luciferase transcriptional reporter fusion. Figure is 3 biological replicates with the shaded region representing the standard deviation.

To study this redundancy, we generated luciferase reporter fusions to look at promoter activity of *canA*, *canB*, and the *ndhFmpsBcanC* complex operon in different growth conditions. We observed low levels of expression during growth in LB. When tested in minimal media glucose (MM-glucose) and malate (MM-malate) we noted a substantial increase from the promoter region that drives expression of the *ndhFmpsBcanC* operon (Fig. 5B-5D). The *canC* gene is separated from the upstream *ndhFmpsB* genes by a 76 bp intergenic region and our expression results suggest that this region may also harbor weak promoter activity.

Since bicarbonate is an important pH buffer, we wanted to test if the expression of the CAs or bicarbonate importer would change under different pH conditions. We grew our reporter fusions in MM-glucose and MM-malate and compared expression in acidic (pH=6) and basic (pH=8) conditions (Fig. S6). Only the *canB* promoter had a significant pH-dependent shift, with notably higher expression in basic vs. acidic pH in MM-glucose, but not in MM-malate. Collectively, these results suggest that the expression of CA and bicarbonate importer genes may be regulated in response to growth conditions, but the underlying mechanisms and the significance of these effects remain to be explored.

### Mutations in *resD* can restore growth of Δ4 with atmospheric levels of CO_2_

Next, we selected for spontaneous suppressor mutations that would allow growth of the Δ4 strain with only atmospheric levels of CO_2_. The Δ4 strain was streaked on LB agar and grown at 37 °C until colonies appeared. After 48 hours, 9 colonies appeared. Upon re-streaking, 5 were found to exhibit detectable growth, at least in regions of high initial cell density. However, only suppressors 1 and 3 were able to form isolated single colonies. Whole genome sequencing revealed that each putative suppressor strain contained one- or two-point mutations (Table 1). The two strains that could grow as single colonies each contained a mutation in *resD*, which encodes a two-component response regulator that controls aerobic and anaerobic respiration. Since ResD affects metabolism in ways that could influence CO_2_ production, we focused on suppressor 1 (carrying a missense mutation in *yppS* and another that changes ResD Arg 201 to Trp) and suppressor 3 (ResD frameshift). In addition, we constructed a derivative of the Δ4 strain that additionally carries a gene deletion, Δ*resD.* Unlike the Δ4 parent strain, all three strains with *resD* mutations grow as isolated colonies and as streaks on LB agar plates (Fig. 6 and S7).

**Figure 6.**
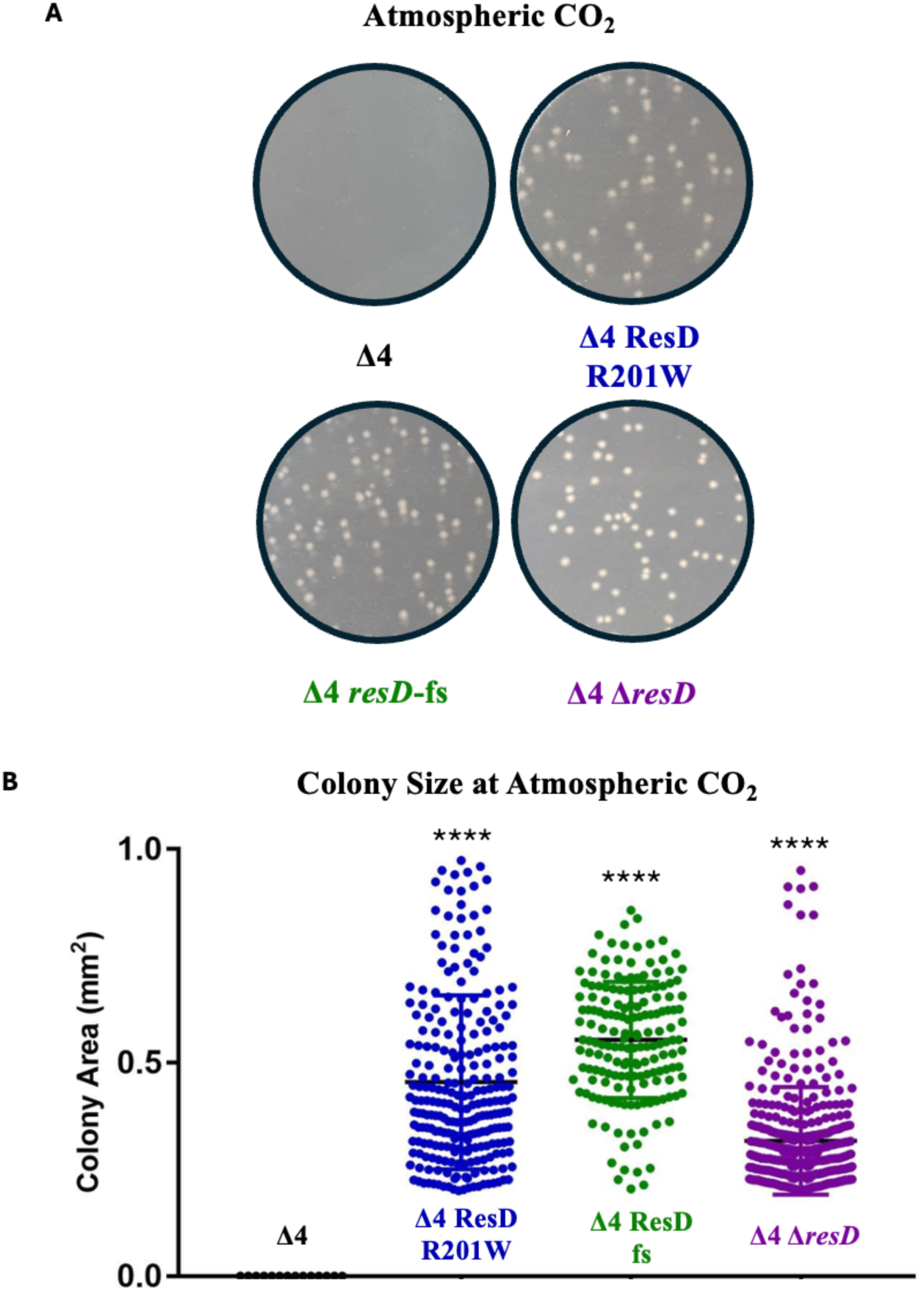
Δ4 strains with mutations in *resD* form colonies at atmospheric CO_2_. a) Well isolated colonies Δ4, Δ4 ResD R201W, Δ4 ResD fs, and Δ4 Δ*resD* were analyzed for size using ImageJ software. Strains were grown on LB agar plates at 37°C at atmospheric CO_2_. Plates were imaged after 19 hours. Figure is representative of 3 independent biological replicates. b) Quantification of colony size of Δ4, Δ4 ResD R201W, Δ4 ResD fs, and Δ4 Δ*resD* at atmospheric CO_2_. P-values were calculated using one-way ANOVA with Tukey’s multiple comparisons test. Statistics are shown for comparison made between mutant with respect to Δ4, **** p<0.0001.

**Table 1.**
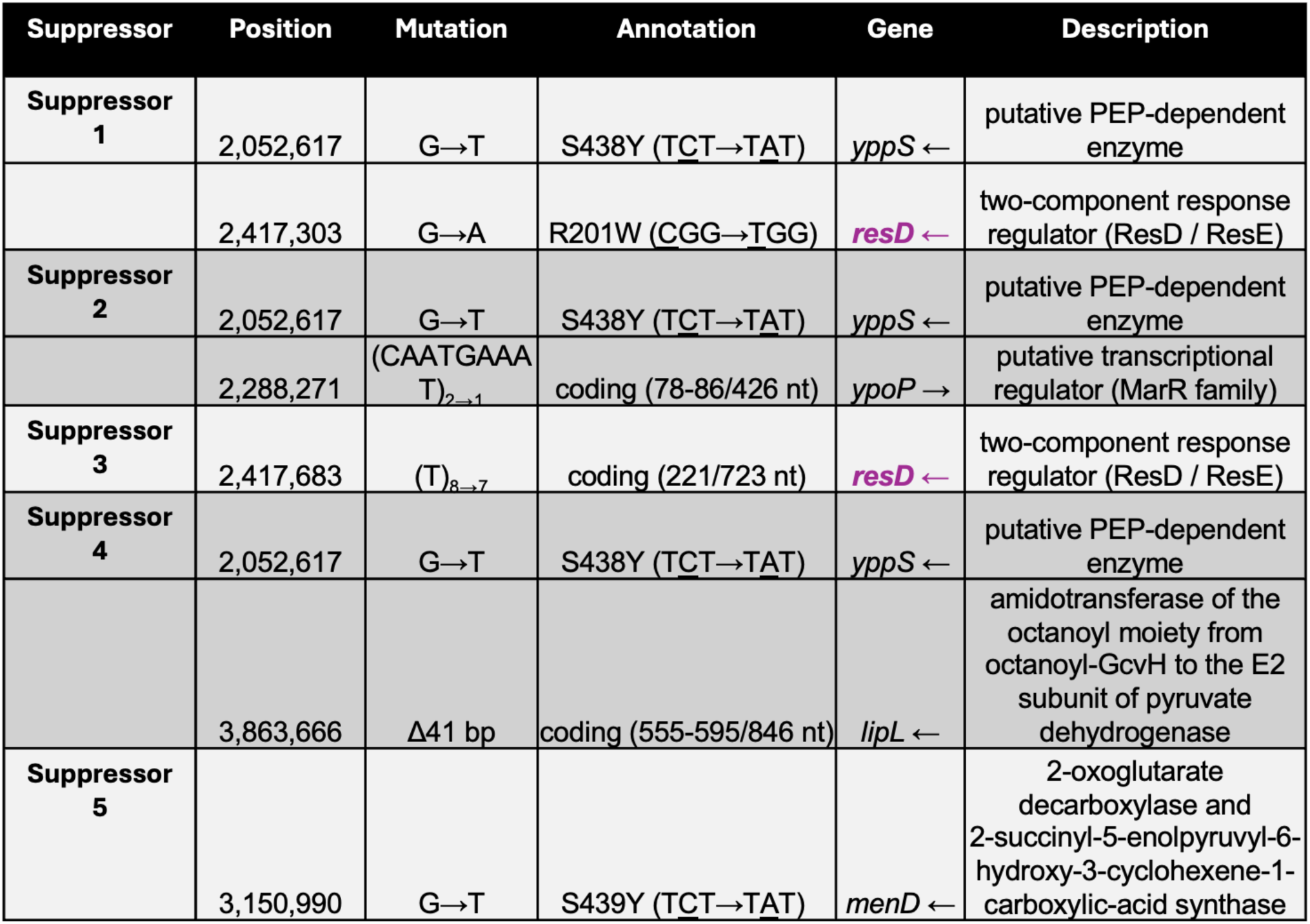
Whole genome sequencing results. List of isolate mutations from adaptive evolution experiment.

Since ResD regulates genes needed for both aerobic and anaerobic respiration (18), we predicted that *resD* mutations could alter the metabolism of the cell to favor CO_2_ production and thereby increase bicarbonate levels. To test this model, we spread lawns of the WT, Δ4, Δ*resD*, and Δ4 Δ*resD* strains on LB medium and incubated at 37 °C for 19 hrs. With a standard size Petri plate incubated in a sealed ∼1.2 L plastic container, we could sample and measure the CO_2_ produced and compare this to the biomass of the resulting lawn (Table S1). Both the WT and Δ4 strain grew well under these conditions with lawns containing comparable amounts of cell mass. We conclude that the Δ4 strain can adapt and ultimately grow under these high cell density conditions, even though it cannot form single colonies. Interestingly, both the *resD* and Δ4 Δ*resD* strains were growth defective, with the mass of the bacterial lawns typically reduced by >5-fold. This suggests that the *resD* mutation significantly reduces cell yield, perhaps be reducing the efficiency of carbon utilization. This model is supported by measurements of net CO_2_ production during lawn growth (Fig. 7, Table S1). Compared to WT (∼1.4 mg CO_2_ released per mg of bacterial lawn), the Δ4 *resD* mutant has ∼3-fold higher CO_2_ production (∼4.4 mg CO_2_ per mg of bacterial lawn). As a result of the lower final biomass in the lawn, the total amount of CO_2_ produced by the Δ4 Δ*resD* strain was lower than either WT or Δ4. Even though the total amount of CO_2_ produced by the Δ4 Δ*resD* strain was lower we infer that the increased localized production was sufficient to support the growth of isolated single colonies.

**Figure 7.**
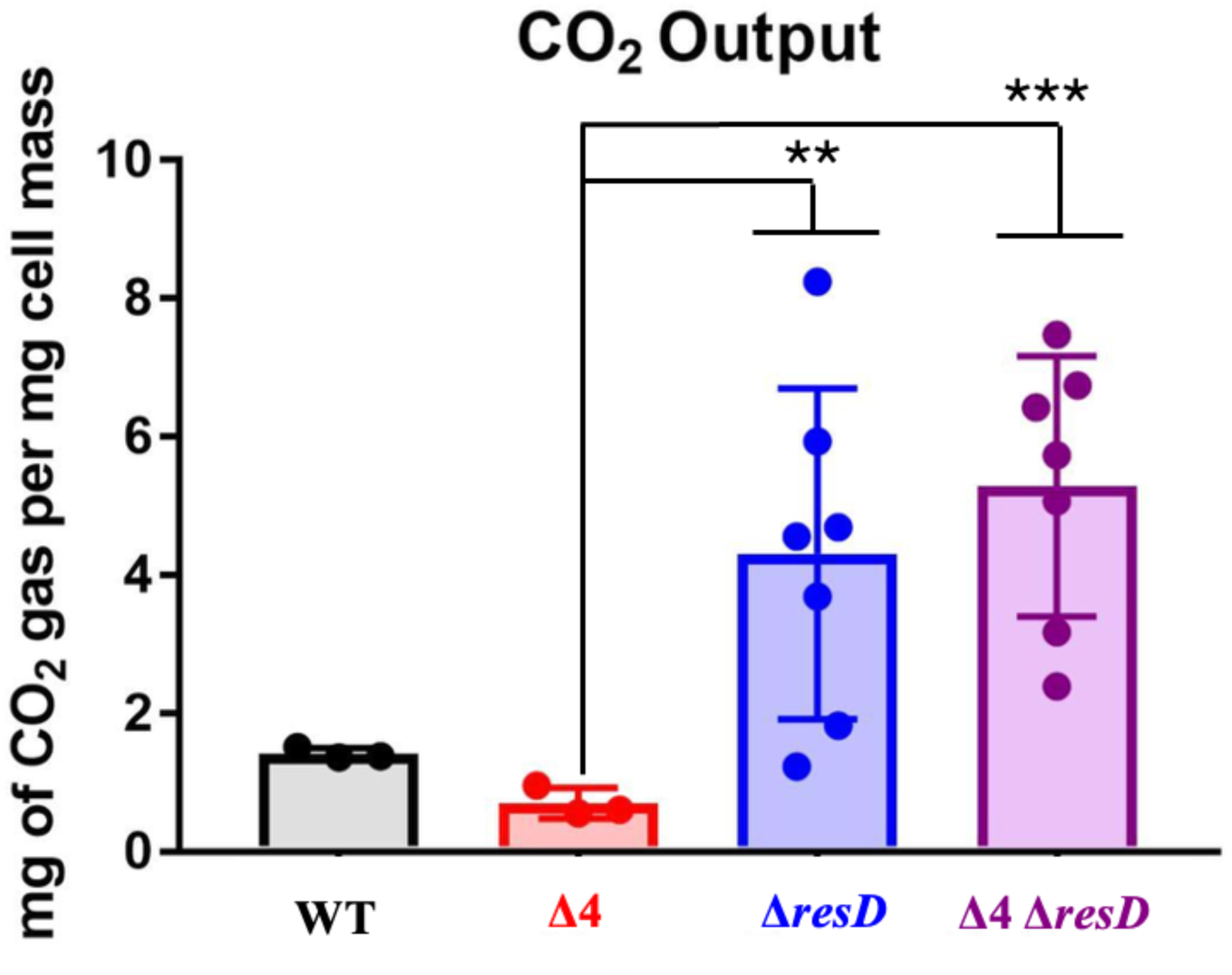
Δ*resD* mutants generate more CO_2_ per milligram of cell mass. Amount of CO_2_ generated by a lawn of WT, Δ4, Δ*resD,* and Δ4 Δ*resD* normalized to cell mass. Each biological replicate has been plotted as a point and significance was analyzed using Welch’s t-test with ** representing p<0.01 and *** representing p<0.001.

## Discussion

Bacteria exhibit remarkably diverse nutritional requirements and complex interlinked metabolic pathways. On one hand, photoautotrophs such as cyanobacteria can meet their carbon and nitrogen needs by fixing atmospheric CO_2_ and N_2_. In contrast, heterotrophic bacteria must obtain carbon and nitrogen through organic molecules. The model organisms *E. coli* and *B. subtilis* are heterotrophs that require fixed sources of both carbon and nitrogen but are otherwise able to grow on simple mineral salts media that provide all other essential elements.

CO_2_ is a relatively low abundance atmospheric gas but is nevertheless required for the growth of many bacteria. The requirement of CO_2_ to support growth of photoautrophs is extensively studied since CO_2_ is the essential substrate for RuBisCo, the first enzyme in the Calvin cycle. CO_2_ is also required for the growth of heterotrophic organisms. It have been known for more than a century that many heterotrophic bacteria are unable to form single colonies when CO_2_ is depleted from air (19). This type of CO_2_ dependence has been periodically rediscovered and revisited over the subsequent decades. The requirement for CO_2_ to support growth is much greater in capnophilic organisms. In studies analogous to those described here for the Δ4 strain, early attempts to cultivate *Brucella* found that its growth was enhanced when cultured in a closed chamber with *B. subtilis* (20). This effect was initially attributed to depletion of O_2_ by *B. subtilis*, however, subsequent work revealed that enhanced growth was instead due to CO_2_ released by *B. subtilis* (20). For most heterotrophs it is not CO_2_ that is growth-limiting, but bicarbonate. Whereas catabolic processes predominantly generate CO_2_, essential anabolic reactions require bicarbonate. The key bicarbonate-dependent enzymes are those for the synthesis of fatty acids, amino acids, and menaquinone precursors (Fig. 8) (6).

**Figure 8.**
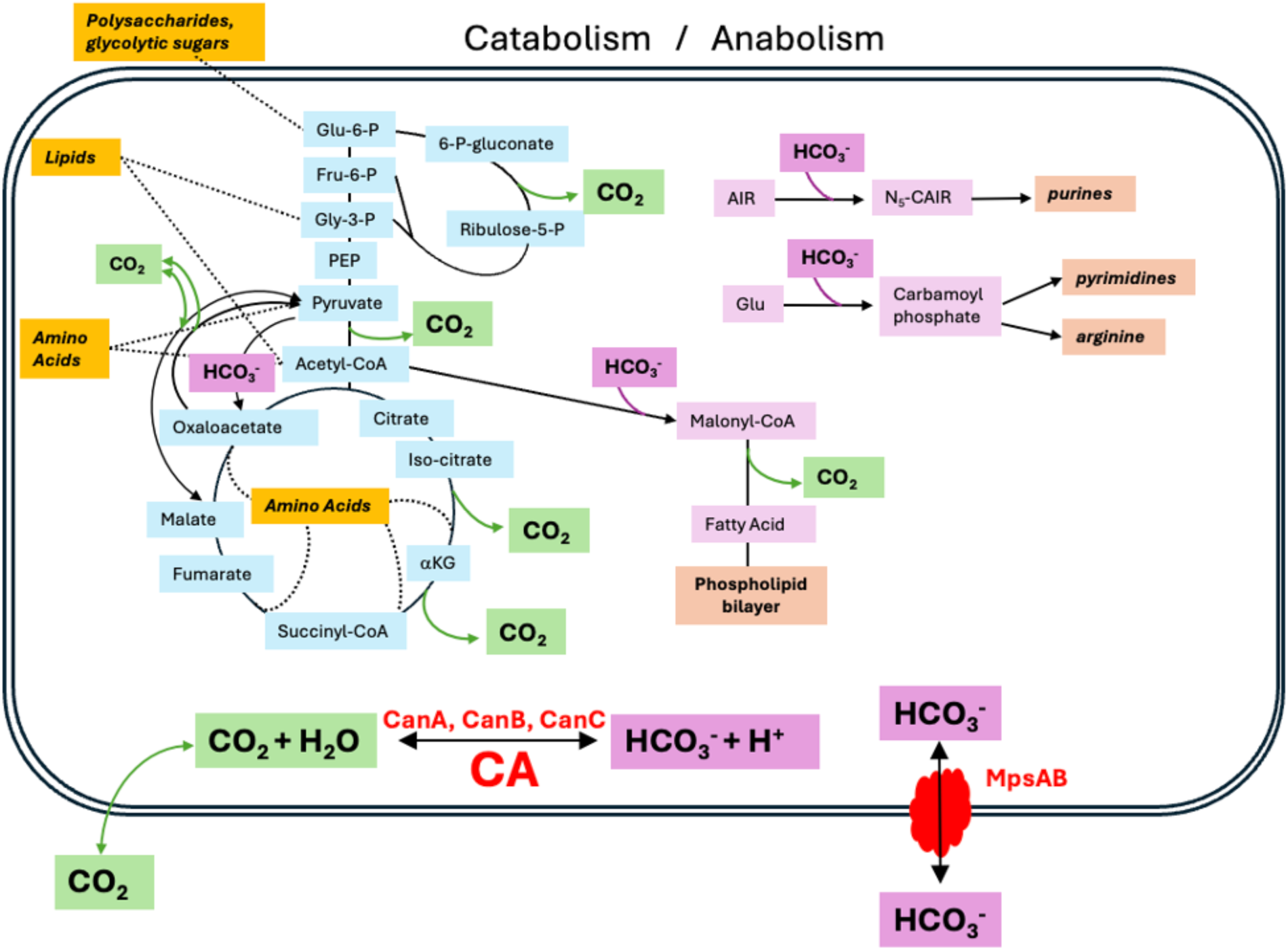
The major roles of CO_2_ and bicarbonate in central metabolism. Major carbon sources (orange) feed into glycolysis and the citric acid cycle (dotted lines). Catabolic reactions (left side) produce CO_2_ (green boxes), with major sources being pyruvate dehydrogenase; isocitrate dehydrogenase and α-ketoglutarate dehydrogenase (citric acid cycle); and 6-phosphogluconate dehydrogenase (pentose phosphate pathway). The malic enzymes that interconvert malate and pyruvate may have a smaller role in CO_2_ production and possibly consumption. CO_2_ can diffuse in and out of the cell or be equilibrated with bicarbonate (HCO_3_^−^) by hydration catalyzed by carbonic anhydrases (CanA, CanB, or CanC; red). Bicarbonate is an important enzyme for anabolic reactions (right side; products in tan boxes), including those that synthesize fatty acids (acetyl-CoA carboxylase), purines (N⁵-CAIR synthetase; PurK), and arginine and pyrimidines (carbamoyl phosphate synthase). Bicarbonate can also be imported by a Na^+^-coupled importer (MpsAB; red).

Quantitative analysis of *E. coli* metabolism suggests that inorganic carbon levels are ∼1000-fold too low to support growth when accounting for CO_2_ loss by diffusion (6). The intracellular level of bicarbonate required to support rapid bacterial growth has not been well defined. Bicarbonate levels are not explicitly included in most metabolic models, although a default estimate of up to 10 mM is sometimes used based solely on values from mammalian physiology. To gain some insight into the likely levels of intracellular bicarbonate needed to support growth we can consider key bicarbonate-dependent enzymes. In general, most of the enzymes in central metabolism have a K_M_ for substrates that is near or even below the atmospheric substrate levels (21). Most of the measured K_M_(HCO_3_^−^) values for bicarbonate-dependent enzymes are in the range of 1 to >10 mM (22, 23). If these K_M_ values are a reliable indication of intracellular concentration, the cell must have mechanisms to increase intracellular bicarbonate levels.

Cells can satisfy their bicarbonate requirement in at least two ways. First, bicarbonate can be concentrated in the cell by active transport (e.g. MpsAB in *S. aureus*) (7). Second, cells can generate sufficient intracellular CO_2_ through metabolism. The latter mechanism is problematic, however, since CO_2_ can escape from cells by diffusion and the spontaneous hydration of CO_2_ to carbonic acid is comparatively slow. This, then, is the key role for CA in heterotrophs. CA can rapidly equilibrate CO_2_ with carbonic acid, and hence bicarbonate to support anabolic enzymes. In addition, the loss of CO_2_ from cells by diffusion may be limited in conditions of high cell density, such as found in biofilms (24, 25).

It is common for organisms to have multiple carbonic anhydrases in their genome, for example, *E. coli* has two carbonic anhydrases, *can* and *cynT*. Can is expressed constituently, while CynT is only expressed during cyanate metabolism (6). In addition, *E. coli* encodes three γ-class CA enzymes (CaiE, PaaY, and YrdA) that are inducible under specific growth conditions (6). Here, we provide physiological evidence that *B. subtilis* encodes three, partially redundant CA enzymes as well as a bicarbonate importer, MpsAB (Fig. 2 and 3). It is unclear why *B. subtilis* has evolved three carbonic anhydrases. One possibility is that they function under different nutritional conditions, developmental states, or in response to specific stresses.

In *B. subtilis*, carbonic anhydrases *canA*, *canB* and *canC* are relatively evenly expressed in LB, MM-glucose and MM-malate (Figs. 5, S6). This suggests that unlike *E. coli*, the redundancy of the CAs is not solely due to carbon source utilization. Alternatively, it is possible that these carbonic anhydrases differ in their preferred metal cofactor. Typically, carbonic anhydrase is a zinc metalloenzyme, however, some microbes have evolved isozymes that rely on other divalent ions, including γ-CA that can use Fe(II) and the zeta-class enzymes (ζ-CA) that use Cd(II). In addition, binding of non-native metal ions to CA often leads to enzyme activity (26, 27). Having CA homologs that use different metals could provide *B. subtilis* with an evolutionary advantage when faced with metal limitation.

It is also possible that cells that lack CA may achieve sufficient intracellular levels of bicarbonate by increasing the metabolic production of CO_2_. Indeed, it has been noted previously that *E. coli* CA-deficient strains can give rise to suppressors that can restore growth in unamended medium, but the causal mutations were not identified (6). Here, we selected for suppressors of Δ4 that could grow at atmospheric CO_2_ and obtained suppressor mutations in *resD*. ResD is an oxygen sensing response regulator of anaerobic and aerobic respiration in *B. subtilis* (28). ResD activate key metabolic genes in *B. subtilis* including *ctaA*, *ctaBCDEFG*, *cydABCD*, *qcrABC,* and *fnr* operons (29). These genes are important for synthesis of heme, cytochrome *caa*_3_, cytochrome *bd*, cytochrome *bc*_1_ and, through Fnr, nitrate reductase (29). Thus, the loss of function of *resD* leads to deficiencies in energy generation. As a result of deficiencies in respiration, cells rely more heavily on substrate level phosphorylation, which reduces metabolic efficiency and increases waste products, including CO_2_. Consistent with this model, we demonstrate that *resD* mutant cells release more CO_2_ per mg of biomass (Fig. 7, Table S1). We therefore suggest that mutations in *resD* restore the growth of Δ4 by generating more endogenous CO_2_ and thus favoring the spontaneous generation of bicarbonate to support cell growth.

In microbial ecology, carbon utilization efficiency (CUE) is often measured using stable isotopes to infer the fraction of metabolized carbon (C) that is incorporated into biomass. Here, we provide a rough estimate of CUE by dividing the biomass C by the total metabolized C that ends up in either biomass or CO_2_ (this ignores the fraction of C that may end up in acetate or other secreted end-products). We find that a *resD* mutation reduces CUE by ∼2-3-fold (Table S1) and therefore infer that the increased rate of localized CO_2_ production is sufficient to support the growth of isolated single colonies.

We conclude that *B. subtilis* employs both CA homologs and a likely bicarbonate importer as bicarbonate concentrating systems. These systems are critical for growth of single colonies, consistent with historical findings (19), but can be bypassed at high cell densities or by increasing atmospheric CO_2_ levels. The isolation of *resD* as a suppressor further highlights the ability of catabolic processes to provide sufficient CO_2_ under conditions where metabolism is altered. Thus, cells can meet much (or all) of their bicarbonate requirement using metabolically generated CO_2_. Remarkably, an analogous situation applies to water: up to 70% of the intracellular water in *E. coli* is generated metabolically (30). Atmospheric CO_2_ levels are currently 50% higher than pre-industrial levels and indoor environments frequently have CO_2_ levels exceeding 1000 ppm (0.1%). As CO_2_ levels rise, this may increase the survival and persistence of bacteria, including pathogens, further adding to the human health risks of elevated CO_2_ (31).

## Materials and Methods

### Strains and growth conditions

Single gene deletions were created using the BKE and BKK collection obtained from the *Bacillus* Genetic Stock Center (BGSC) (32). Single deletions were made clean with pDR244 plasmid and confirmed using check primers (32). Double deletions were made by moving in a marked gene into the desired clean single background. These mutations were selected for on rich medium with proper antibiotic. The rich medium for growth was lysogeny broth (LB) (Affymetrix) with erythromycin (1 μg/mL), lincomycin (25 μg/mL), or kanamycin (15 μg/mL) as needed. All strains used in this study are referenced in Table S2. Double deletions were made clean (32), and a single marked gene was moved to construct triple deletions. Each triple mutant was then made clean. Congression was checked for at every step by PCR amplification using a gene specific upstream and downstream primer listed in Table S3.

To delete two genes in the same operon (*mpsB* and *canC*), long flanking homology PCR was used. Using the gDNA of erythromycin marked *mpsB* and *canC* as a template, 200 base pairs upstream of *mpsB* and 200 base pairs downstream of *canC* was amplified along with the erythromycin cassette. These two PCR products were then stitched together using the homology of the erythromycin cassette. This final PCR product was then transformed into WT *B. subtilis* before Δ*mpsBcanC*::*erm* gDNA was extracted and moved into desired genetic backgrounds. For construction of Δ4, marked Δ*mpsBcanC*::*erm* gDNA was transformed into clean Δ*canA* Δ*canB* background. Strains were selected for on proper antibiotic and congression was checked for by PCR.

### Revival from cryogenic stock

For plate images, WT and mutant strains were streaked from frozen glycerol stocks onto LB agar plates (Affymetrix). Plates were grown overnight at 37 °C at atmospheric, 0.5%, or 5% CO_2._ To generate 5% CO_2_ conditions, a 1.2 L container was allowed to equilibrate in a 5% CO_2_ incubator for at least 1 hr. Similarly, for ∼0.5% CO_2_ three 50 mL Falcon tubes were allowed to equilibrate at 5% CO_2_ before being added to a 1.2 L container. After 1 hr., the plates were added to the container, sealed, and placed in a 37 °C incubator overnight. Plates were imaged at 19 hours. All images were taken at the same magnification and are consistent throughout the entire paper.

### Bioinformatics

Proteins with homology to CanA were identified using HHPred (MPI) (33). Weak similarity was noted to the hypothetical protein *mpsB*, which is annotated as a possible bicarbonate transporter. Sequence alignments were generated using MUSCLE (EMBL server) (34). Structural alignments were made using the matchmaker command in ChimeraX (35) with the AlphaFold predicted protein structure (36).

### Transcriptional luciferase fusions

The promoter of *canA*, *canB*, *canC*, and *mpsB* were cloned by restriction digestion and subsequent ligation into pBS3K-lux plasmid. These plasmids were then transformed into WT *B. subtilis* and selected for on LB with appropriate antibiotic erythromycin (1 μg/mL) or kanamycin (15 μg/mL). Strains containing this plasmid were then grown in LB, MM-Glucose (pH 6, 7.4, or 8), or MM-Malate (pH 6, 7.4, or 8). Chemically defined minimal medium (MM) contained 10 g/L ammonium sulfate (NH_4_)_2_SO_4_, 5 g/L trisodium citrate (Na_3_C_6_H_5_O_7_•2H_2_O), 5 g/L l-glutamic acid (potassium salt monohydrate), 40 mM MOPS pH 7.4 for neutral pH, MOPS pH 6 for acidic media, and MOPS pH 8 for basic media, 2 mM KPO_4_ (pH 7.0), 10 mg/L tryptophan, 0.8 mM MgSO_4_, 50 μM ferric ammonium citrate, 5 μM MnCl_2_, and either 0.8% (w/v) malate pH 7.4 (MM-malate) or 2% (w/v) glucose (MM-glucose) as carbon source. Growth was monitored in a plate reader at 37 °C, shaking, for 24 hours. The relative luminescence (RLU) was normalized to growth (OD_600_).

### Determination of colony size

For growth under various initial CO_2_ concentrations, we used air-tight plastic food storage containers (∼12 x 12 x 8 cm; maximal vol. of 1216 mL as measured by weight of water when full) with a silicone gasket and plastic snap closures. Empty containers were found to maintain elevated CO_2_ concentrations for >19 hrs with no detectable loss. All bacterial strains were streaked from frozen glycerol stocks onto LB agar plates and grown overnight at 37 °C at atmospheric CO_2_. For strains with atmospheric growth defects, strains were streaked from frozen glycerol stocks onto LB agar plates and grown overnight at 5% CO_2_. Liquid suspension of OD_600_ ∼0.1 was made from strains grown overnight. 100 µL of 10^−5^ and 10^−6^ serial diluted cells were plated on LB agar plates (20 mL) and placed in the container. For the lawn of WT for co-incubation, an inoculum of WT was grown to OD_600_ = 0.4 and 100 µL of this culture was spread onto a LB agar plate and placed in a container with the serial diluted mutant plates. Plates were incubated at 37 °C overnight and imaged at 19 hrs. Well-isolated colonies were analyzed using Fiji-ImageJ software to determine the area of the colonies (37).

### Suppressor selection

The Δ4 parent stain was streaked from frozen glycerol stock onto a LB agar plate and grown at 37 °C at atmospheric CO_2_ until isolates appeared (48 hours). Each isolate was then grown shaking overnight at 37 °C at atmospheric CO_2_. From these overnight cultures, glycerol stocks were prepared and isolates were frozen at −80 °C overnight. From frozen glycerol stocks, the isolates were re-streaked on fresh LB agar plates and grown overnight at 37 °C at atmospheric CO_2_. Genomic DNA was extracted using a kit (ThermoFisher) from isolates that were able to grow overnight in these unfavorable conditions. The isolated genomic DNA was sent for whole genome sequencing (SeqCenter) along with the Δ4 parent strain. Single nucleotide variations were analyzed by SeqCenter by comparing our extracted DNA to 168 (NC_000964.3) *B. subtilis* genome. Individual isolate differences from the Δ4 parent strain were determined by comparing the list of point mutations in our isolate genome to Δ4.

### Measurement of CO_2_ output

WT, Δ4, Δ*resD*, Δ4 Δ*resD* were grown fresh from frozen glycerol stock overnight at 37 °C on LB agar plates. All strains, except for Δ4 were grown at atmospheric CO_2_. From the LB plate a liquid suspension of OD_600_ = 0.4 was made. To make a lawn, 100 µL of these cells were spread with beads on a fresh LB agar plate and incubated in a sealed 1.2 L plastic container to monitor CO_2_ production. Concentrations of CO_2_ were measured using a Temtop C1 portable indoor air quality monitor, which was chosen for its small size (8.9 x 6.5 x 1.8 cm) to fit within the sealed plastic containers. These monitors function over the range of 0.04 to 0.5% with an accuracy of 5% and over a wide range of atmospheric humidity. CO_2_ monitors were calibrated by exposure to outside air as described by the manufacturer prior to each use. Trial experiments revealed that CO_2_ levels in the growth containers exceed the linear range of these monitors. Therefore, we sampled and then diluted the atmospheric gases from each growth chamber prior to measurement. Bacterial lawns were grown for 19 hours at 37 °C in an air-tight container (box 1) with a 50 mL Falcon tube that had the opening sealed with parafilm containing a small (∼5-10 mm^2^) hole to allow gas exchange (measured gas volume = 59 mL). After incubation, the Falcon tubes were moved to a fresh container (box 2) with a CO_2_ monitor and sealed. CO_2_ readings were recorded prior to introduction of the Falcon tube (starting CO_2_) and the final CO_2_ after 45 min. equilibration (final CO_2_). Using the starting and final CO_2_ readings, and the gas volume of box 1 and box 2 (adjusted for air displaced by plasticware and the CO_2_ monitor), we calculated the net CO_2_ produced during lawn growth (Table S1). The amount of CO_2_ generated was normalized against the wet weight of the corresponding bacterial lawn.

## Acknowledgements

This work was supported by the National Institute of Health awards R35GM122461 (to J. D. H.). The content is solely the responsibility of the authors and does not necessarily represent the official views of the National Institutes of Health.

## Supplementary Material

**Table S1.** Production of CO_2_ gas during growth of bacterial lawns.

| Strain | Lawn Weight box 1 (mg) | Starting CO<sub>2</sub> box 2 (ppm) | Final CO<sub>2</sub> box 2 (ppm) | Final CO<sub>2</sub> box 1 (%) | CO<sub>2</sub> in box 1 (net mg) | CO<sub>2</sub> gas per bacterial lawn (mg/mg) | Estimated CUE (%) |
| --- | --- | --- | --- | --- | --- | --- | --- |
| WT #1 | 70.00 | 500 | 2848 | 4.50 | 95.2 | 1.36 | 21.2 |
| WT #2 | 70.00 | 440 | 2829 | 4.57 | 96.8 | 1.38 | 20.9 |
| WT #3 | 70.00 | 449 | 3070 | 5.01 | 106.2 | 1.52 | 19.5 |
| Δ4 #1 | 60.00 | 500 | 1911 | 2.72 | 57.2 | 0.95 | 27.8 |
| Δ4 #2 | 70.00 | 510 | 1549 | 2.02 | 42.2 | 0.60 | 37.8 |
| Δ4 #2 | 70.00 | 460 | 1405 | 1.84 | 38.2 | 0.55 | 40.1 |
| Δ*resD* #1 | 20.00 | 500 | 1398 | 1.75 | 36.4 | 1.82 | 16.7 |
| Δ*resD* #2 | 30.00 | 560 | 1469 | 1.78 | 37.0 | 1.23 | 22.9 |
| Δ*resD* #3 | 5.00 | 443 | 1177 | 1.43 | 29.6 | 5.93 | 5.8 |
| Δ*resD* #3 | 7.30 | 430 | 1253 | 1.60 | 33.2 | 4.55 | 7.4 |
| Δ*resD* #4 | 5.50 | 426 | 1548 | 2.17 | 45.3 | 8.24 | 4.3 |
| Δ*resD* #5 | 6.40 | 484 | 1067 | 1.15 | 23.6 | 3.69 | 9.0 |
| Δ*resD* #6 | 5.40 | 475 | 1101 | 1.23 | 25.3 | 4.69 | 7.2 |
| Δ4 *resD* #1 | 3.00 | 500 | 975 | 0.95 | 19.3 | 6.42 | 5.4 |
| Δ4 *resD* #2 | 20.00 | 515 | 1067 | 1.10 | 22.4 | 1.12 | 24.6 |
| Δ4 *resD* #3 | 5.00 | 400 | 1236 | 1.62 | 33.7 | 6.74 | 5.2 |
| Δ4 *resD* #4 | 7.30 | 439 | 1474 | 2.00 | 41.8 | 5.73 | 6.0 |
| Δ4 *resD* #5 | 8.50 | 443 | 1506 | 2.06 | 43.0 | 5.06 | 6.8 |
| Δ4 *resD* #6 | 9.50 | 459 | 1022 | 1.11 | 22.7 | 2.39 | 13.3 |
| Δ4 *resD* #7 | 8.60 | 496 | 1168 | 1.32 | 27.2 | 3.17 | 10.4 |

**Table S2:** Strain List.

| Strain | Genotype | Construction | Reference |
| --- | --- | --- | --- |
| CU1065 | WT (*trpC2*) | Lab strain | Lab stock |
| HB31088 | $\Delta canC::erm$ | BGSC gDNA→CU1065 | This work |
| HB31091 | $\Delta canB::erm$ | BGSC gDNA→CU1065 | This work |
| HB31094 | $\Delta canB$ | pDR244→HB31091 | This work |
| HB31099 | $\Delta canC$ | pDR244→HB31088 | This work |
| HB31105 | $\Delta canA::erm$ | BGSC gDNA→CU1065 | This work |
| HB31108 | $\Delta canA::erm \Delta canB$ | HB31105 gDNA→HB31094 | This work |
| HB31113 | $\Delta canA::erm \Delta canC$ | HB31105 gDNA→HB31088 | This work |
| HB31118 | $\Delta canB::erm \Delta mpsB$ | HB31091 gDNA→HB31088 | This work |
| HB31123 | $\Delta canA$ | pDR244→HB31105 | This work |
| HB31126 | $\Delta canA \Delta canB$ | pDR244→HB31108 | This work |
| HB31131 | $\Delta canA \Delta canC$ | pDR244→HB31113 | This work |
| HB31136 | $\Delta canB \Delta canC$ | pDR244→HB31118 | This work |
| HB31141 | $\Delta canA \Delta canB$<br>$\Delta canC::erm$ | HB31088 gDNA→HB31126 | This work |
| HB31191 | $\Delta mpsB::erm$ | BGSC gDNA→CU1065 | This work |
| HB31219 | $\Delta mpsB canC::erm$ | PCR Fusion HB31191 +<br>HB31088→CU1065 | This work |
| HB31231 | $\Delta canA \Delta mpsB::erm$ | HB31191 gDNA→HB31123 | This work |
| HB31232 | $\Delta canA$<br>$\Delta mpsB canC::erm$ | HB31219 gDNA→HB31123 | This work |
| HB31233 | $\Delta canB$<br>$\Delta mpsB canC::erm$ | HB31219 gDNA→HB31094 | This work |
| HB31234 | $\Delta canA \Delta canB$<br>$\Delta mpsB canC::erm (\Delta 4)$ | HB31219 gDNA→HB31126 | This work |
| HB31237 | $\Delta canA \Delta canB$<br>$\Delta mpsB::erm$ | HB31191 gDNA→HB31126 | This work |
| HB31239 | $\Delta canA \Delta canB \Delta mpsB$ | pDR244→HB31237 | This work |
| HB31240 | $\Delta canA \Delta mpsB canC$ | pDR244→HB31232 | This work |
| HB31243 | $\Delta canB \Delta mpsB canC$ | pDR244→HB31233 | This work |
| HB31666 | $\Delta mpsB::kan$ | BGSC gDNA→CU1065 | This work |
| HB31479 | CU1065 $sacA::P_{mpsB}$-<br>$luxABCDE (erm)$ | Promoter-$mpsB$-<br>pBS3Elux→CU1065 | This work |
| HB31482 | CU1065 $sacA::P_{canC}$-<br>$luxABCDE (erm)$ | Promoter-$canC$-<br>pBS3Elux→CU1065 | This work |
| HB31484 | CU1065 $sacA::P_{canB}$-<br>$luxABCDE (erm)$ | Promoter-$canB$-<br>pBS3Elux→CU1065 | This work |
| HB31490 | $\Delta canB::erm$<br>$\Delta mpsB::kan$ | HB31666 gDNA→HB31091 | This work |
| HB31493 | CU1065 $sacA::P_{canA}$-<br>$luxABCDE (kan)$ | Promoter-$canA$-<br>pBS3Klux→CU1065 | This work |
| HB31499 | $\Delta canA \Delta canB$<br>$\Delta mpsB canC::erm$<br>$resD$ R201W | $\Delta 4$ suppressor 1 | This work |
| HB31501 | $\Delta canA \Delta canB$<br>$\Delta mpsB canC::erm$<br>$resD$ fs | $\Delta 4$ suppressor 3 | This work |
| HB31525 | $\Delta resD::kan$ | BGSC gDNA→CU1065 | This work |
| HB31534 | $\Delta canA \Delta canB$<br>$\Delta resD::kan$ | HB31525→HB31126 | This work |
| HB31528 | $\Delta canA \Delta canB$<br>$\Delta mpsB canC::erm$<br>$\Delta resD::kan$ | HB31219→HB31534 | This work |

**Table S3:**
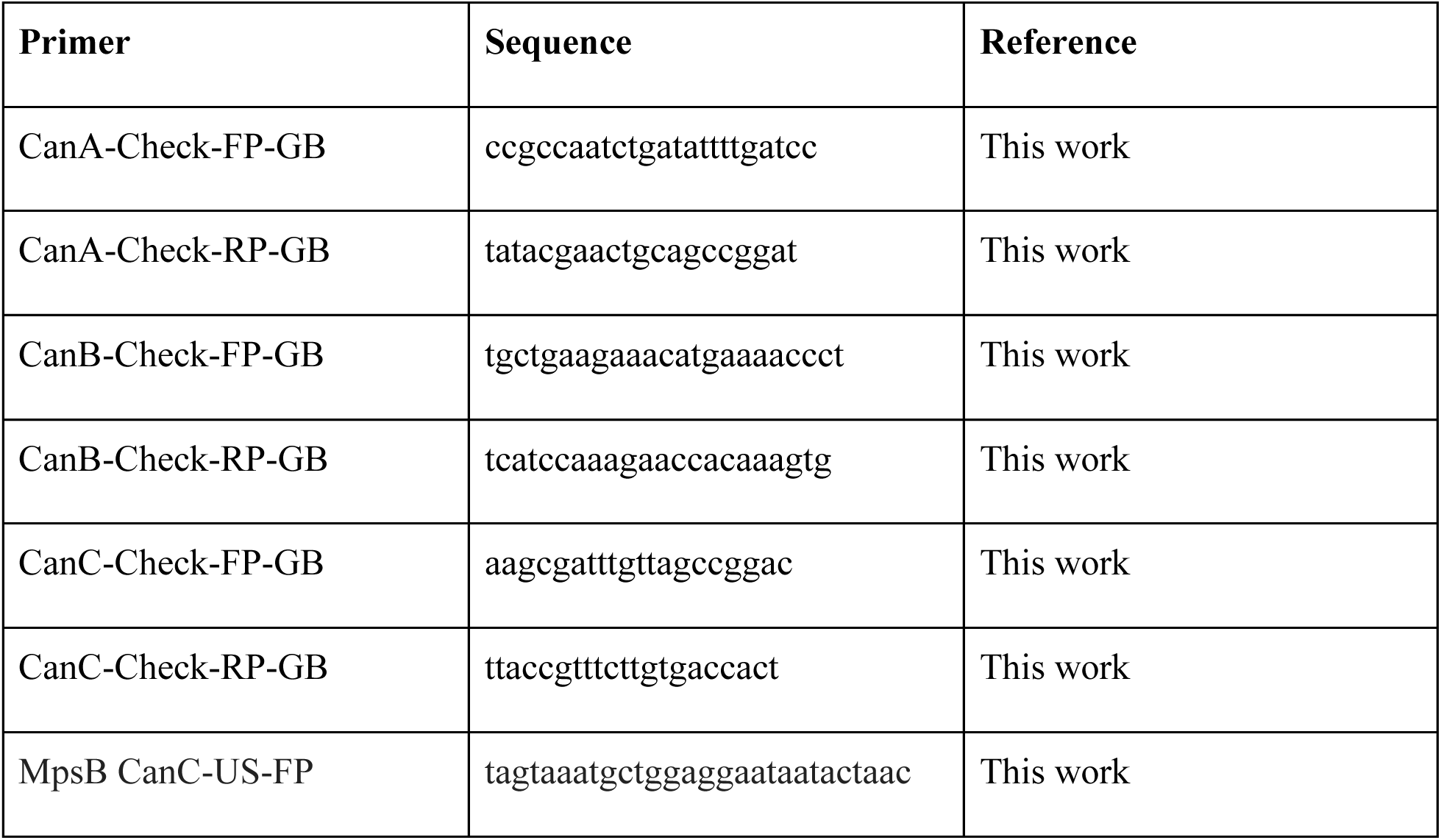
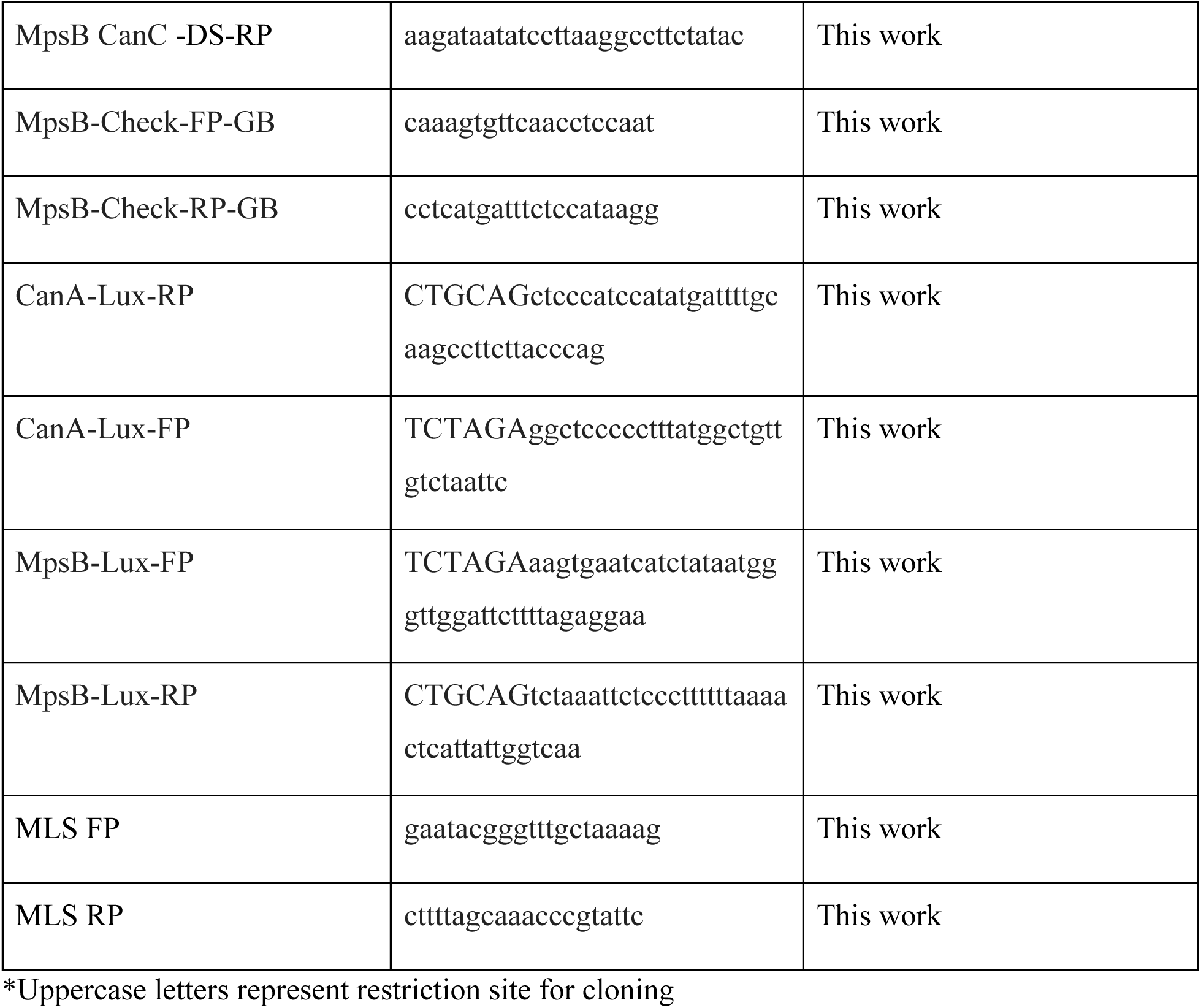
Primer List.

## Supplementary Figures

**Figure S1.**
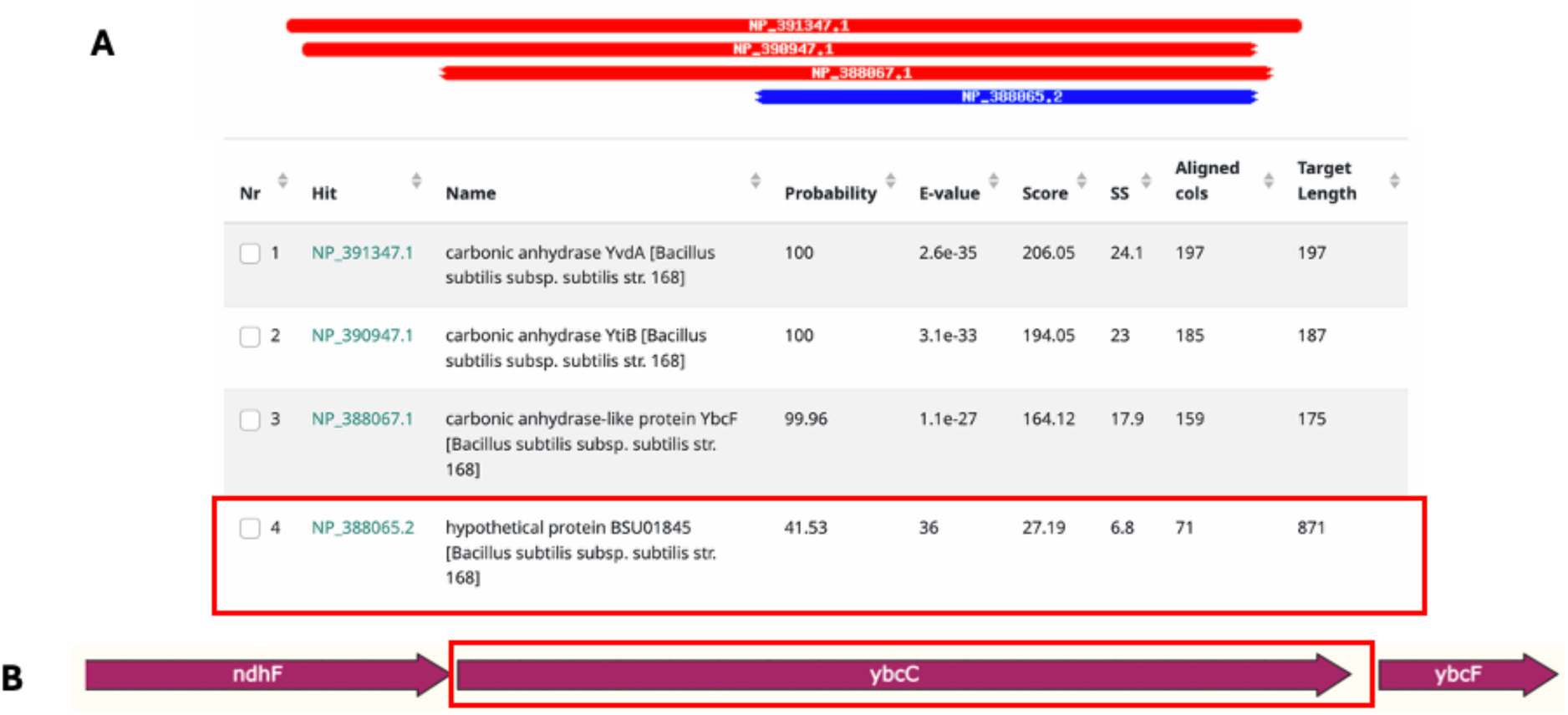
Bioinformatic identification of MpsB (YbcC). a) The protein sequence of CanA (YvdA) was used to search for homologs using HHPred. The expected CAs were identified together with a much more distantly related hypothetical protein, *ybcC*. b) The *ybcC* gene is encoded together with *ndhF.* Together, these two genes encode predicted subunits of a candidate MpsAB-type bicarbonate importer.

**Figure S2.**
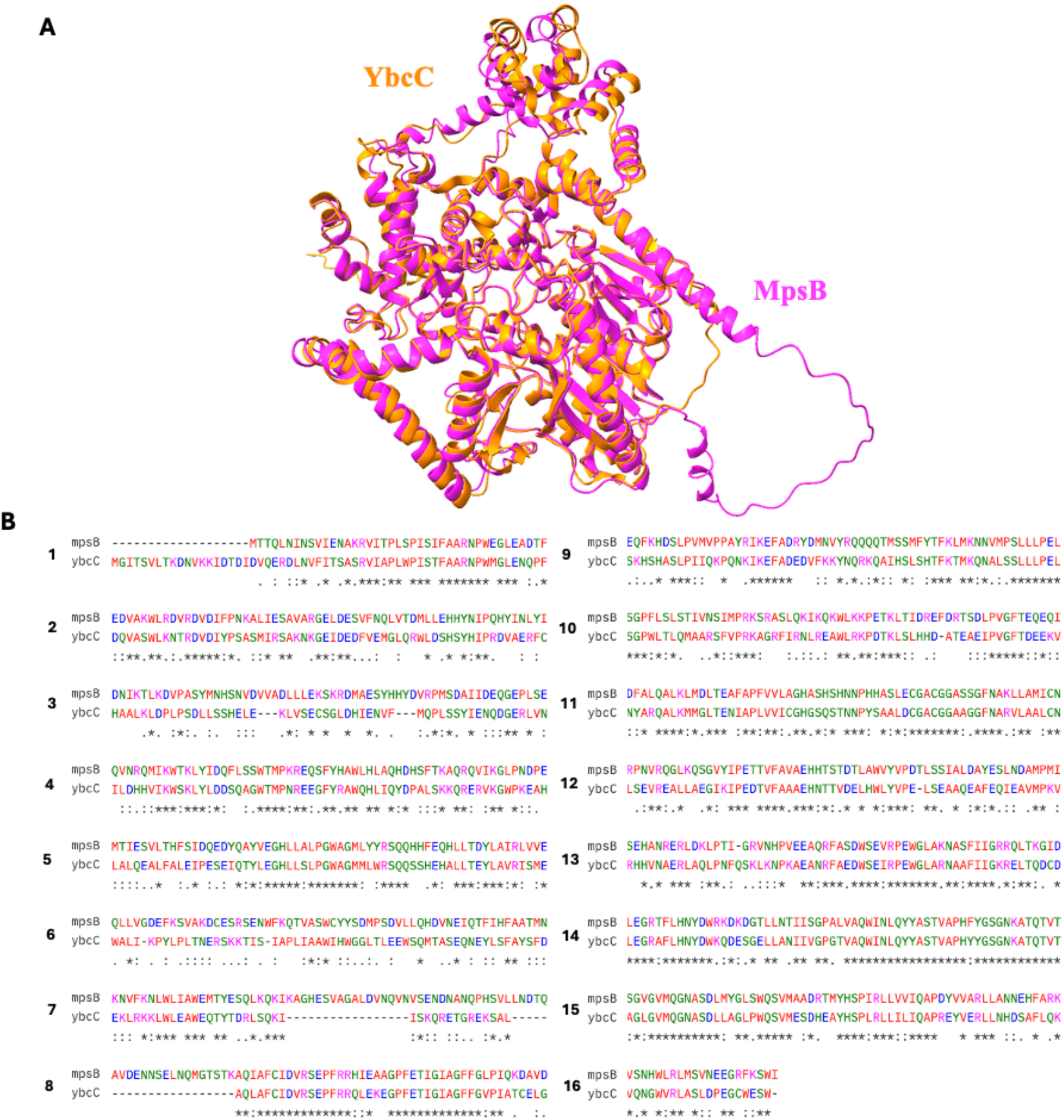
*Bacillus subtilis* MpsB (YbcC) shares structure and sequence similarity to *S. aureus* bicarbonate transporter MpsB. a) Predicted structure of *B. subtilis* YbcC (orange) aligned with *S. aureus* bicarbonate transporter MpsB (pink). Aligned using matchmaker command in ChimeraX. b) Multiple sequence alignment of *B. subtilis* MpsB (YbcC) and *S. aureus* MpsB, aligned with MUSCLE (EMBL server).

**Figure S3.**
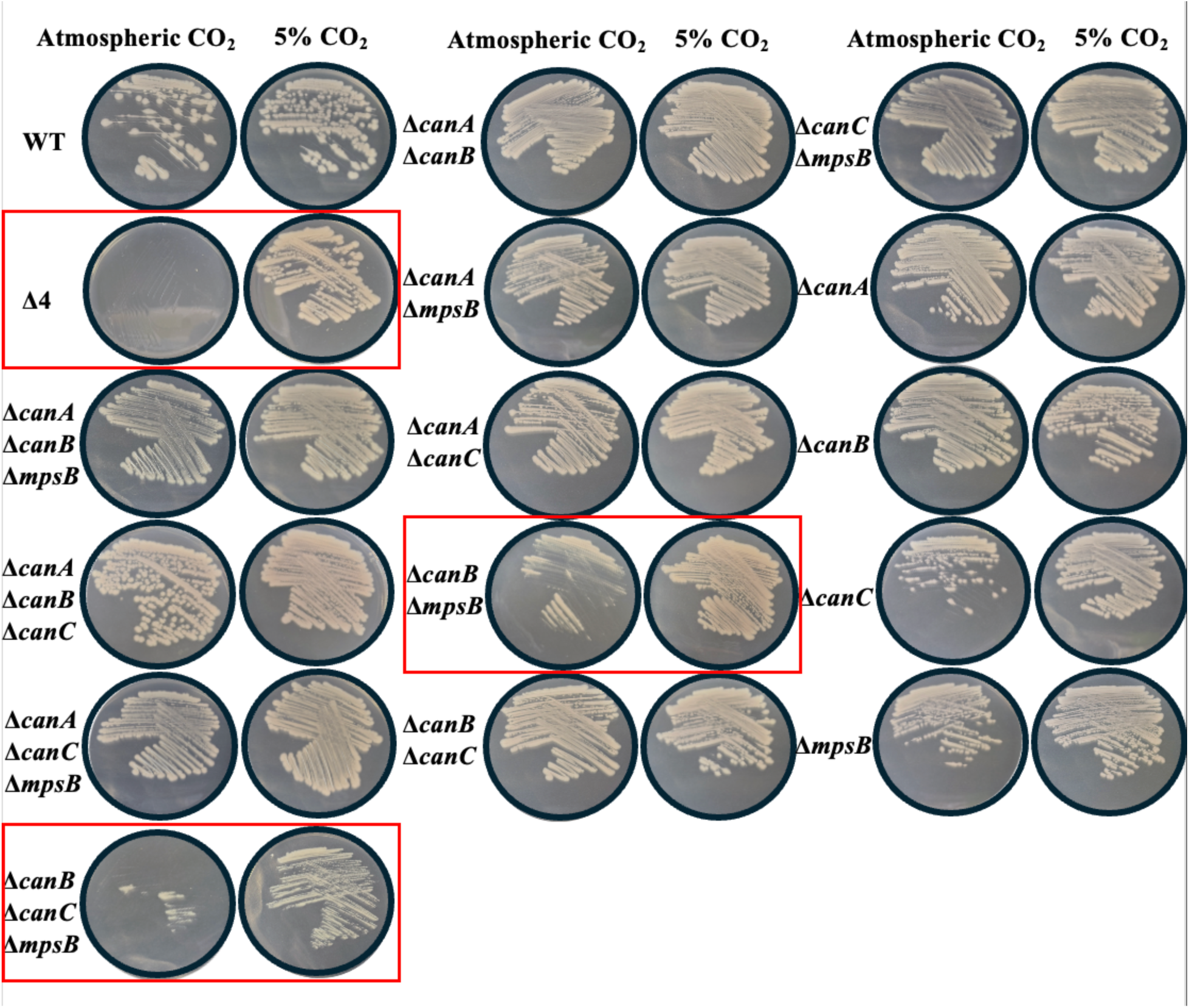
Streaks of all single, double, triple, and quadruple mutants. WT, Δ4, Δ*canA* Δ*canB* Δ*mpsB*, Δ*canA* Δ*canB* Δ*mpsB,* Δ*canA* Δ*mpsBcanC*, Δ*canB* Δ*mpsBcanC*, Δ*canA* Δ*canB*, Δ*canA* Δ*mpsB*, Δ*canA* Δ*canC*, Δ*canB* Δ*mpsB*, Δ*canB* Δ*canC*, Δ*mpsBcanC*, Δ*canA*, Δ*canB*, Δ*canC*, and Δ*mpsB* streaked out on LB agar plates directly from −80° C stocks and then grown at 37°C at atmospheric CO_2_ or 5% CO_2_. Plates were imaged after 19 hours. Figure is representative of 3 independent biological replicates.

**Figure S4.**
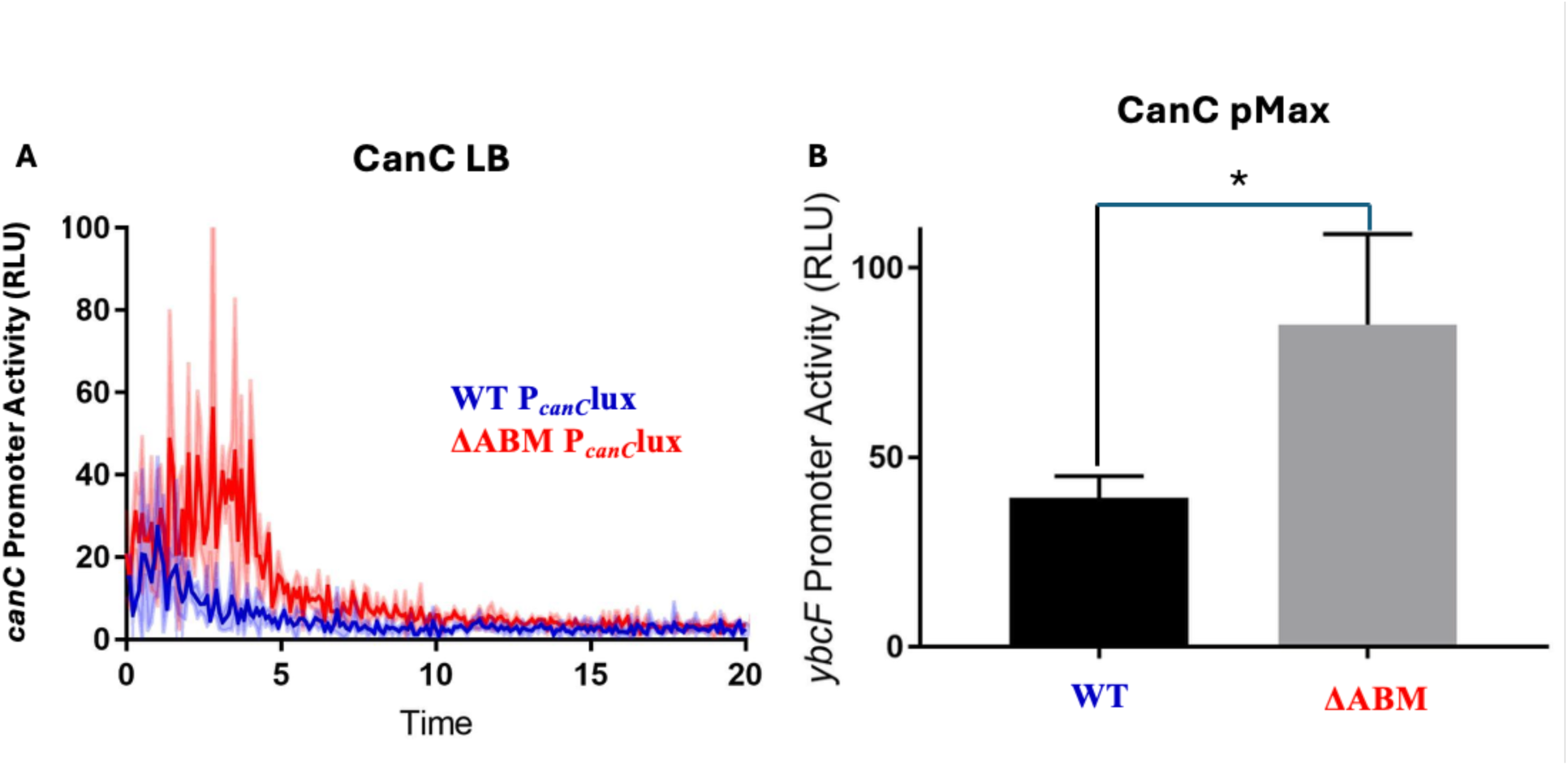
Transcriptional expression of *canC* goes up in a in a ΔABM background. a) Expression of *canC* in WT and Δ*canA* Δ*canB* Δ*mpsB* (ΔABM) background when grown in LB media. Graph is three biological replicates with the shaded area representing standard deviation. b) Bar graph of the pMax of *canC* (WT) and *canC* (ΔABM) in LB media. Figure is three biological replicated with the error bars showing standard deviation. Statistics were calculated using one-way ANOVA with Tukey’s multiple comparisons, * p<0.05.

**Figure S5.**
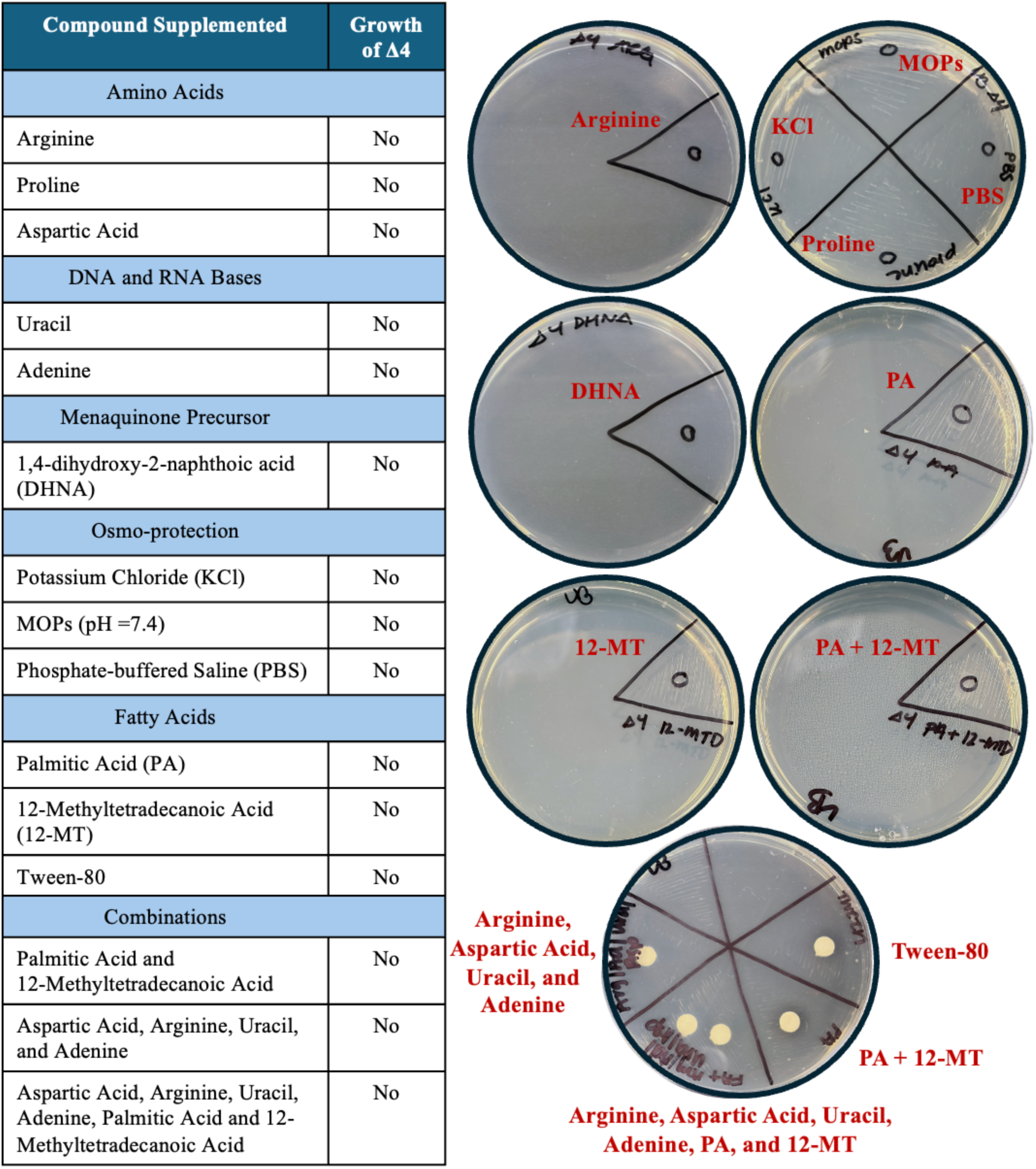
Compounds supplemented to restore the growth of Δ4. a) Table of compounds supplemented and b) images of plates with Δ4 streaked with the supplemented compounds spotted directly on the plate or on filter disk. No compound or combinations or compounds restored the growth of Δ4.

**Figure S6.**
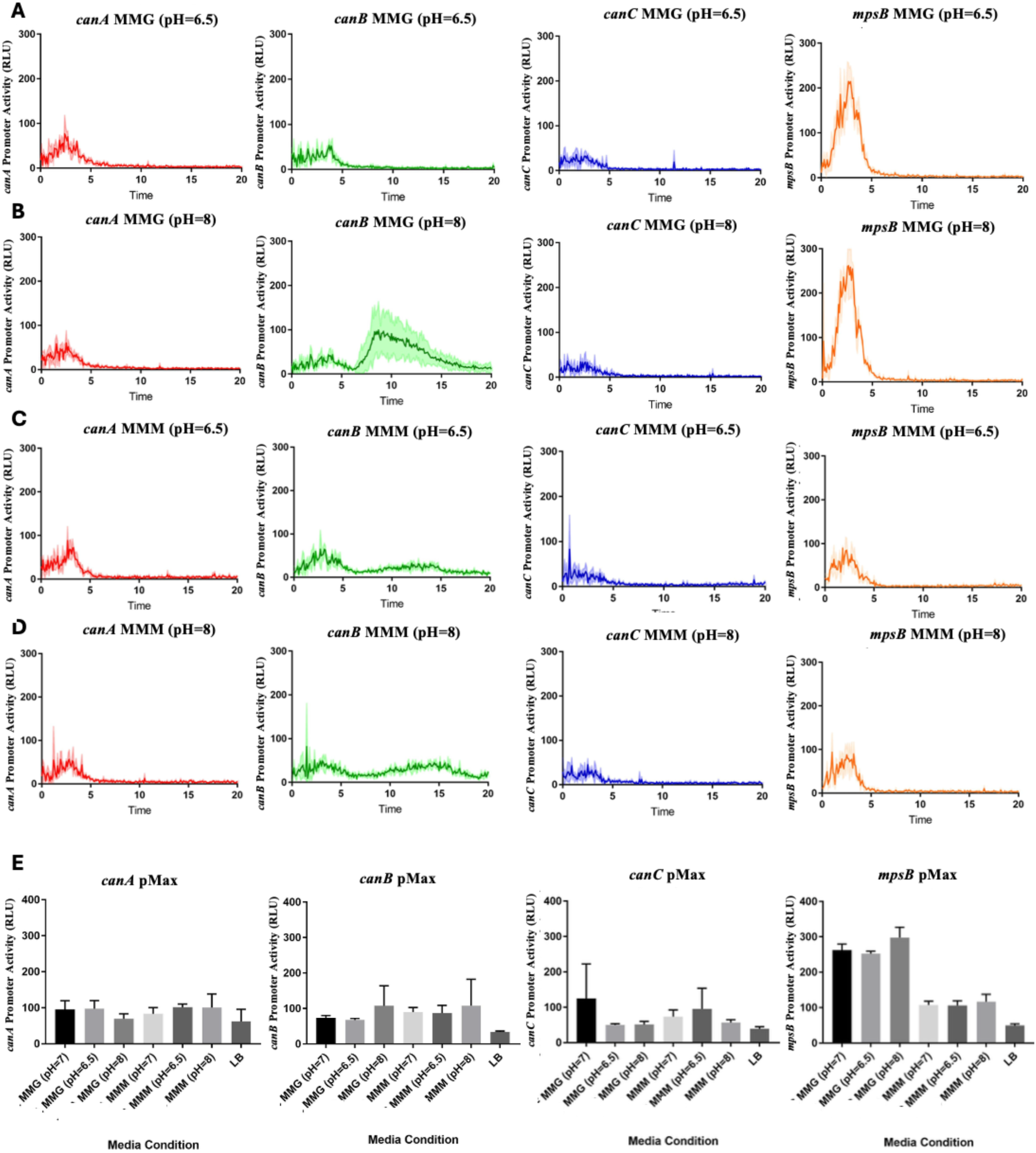
*canB* responds to basic conditions. Expression of *canA*, *canB*, *canC*, and *mpsB* in a) acidic MM-glucose (pH=6.5), b) basic MM-glucose (pH=8), c) acidic MM-malate (pH=6.5), and d) basic MM-Malate (pH=8). Expression monitored using luciferase transcriptional reporter fusion. Figure is 3 biological replicates with the shaded region representing the standard deviation. e) Bar graph of the pMax of *canA*, *canB*, *canC*, and *mpsB* grown at each condition. Figure is 3 biological replicates with the error bar representing the standard deviation.

**Figure S7.**
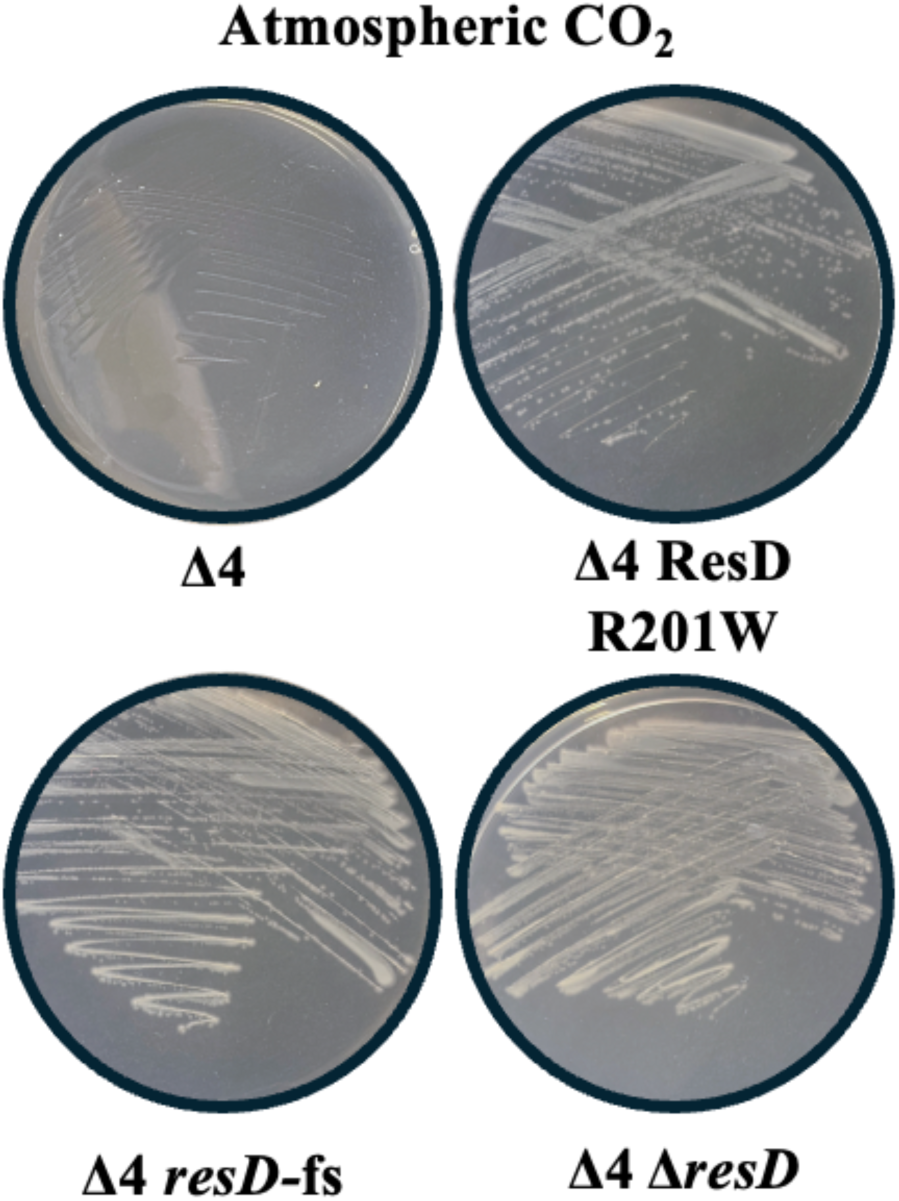
Δ4 with mutations in *resD* grow at atmospheric CO_2_. Δ4, Δ4 ResD R201W, Δ4 ResD fs, and Δ4 Δ*resD* streaked out on LB agar plates grown at 37°C at atmospheric CO_2_. Plates were imaged after 19 hours. Figure is representative of 3 independent biological replicates.

## References

1. Meldrum NU, Roughton FJ. 1933. The state of carbon dioxide in blood. J Physiol 80:143–70.

2. Smith KS, Jakubzick C, Whittam TS, Ferry JG. 1999. Carbonic anhydrase is an ancient enzyme widespread in prokaryotes. Proc Natl Acad Sci U S A 96:15184–9.

3. Franco MEE, Singer E, Roux S, Meredith LK, U’Ren JM. 2026. Genomic and metagenomic survey of microbial carbonic anhydrase genes reveals novel clades, high diversity, and biome specificity. ISME Commun 6:ycag054.

4. Geers C, Gros G. 2000. Carbon dioxide transport and carbonic anhydrase in blood and muscle. Physiol Rev 80:681–715.

5. Badger M. 2003. The roles of carbonic anhydrases in photosynthetic CO(2) concentrating mechanisms. Photosynth Res 77:83–94.

6. Merlin C, Masters M, McAteer S, Coulson A. 2003. Why is carbonic anhydrase essential to *Escherichia coli*? J Bacteriol 185:6415–24.

7. Fan SH, Ebner P, Reichert S, Hertlein T, Zabel S, Lankapalli AK, Nieselt K, Ohlsen K, Gotz F. 2019. MpsAB is important for *Staphylococcus aureus* virulence and growth at atmospheric CO(2) levels. Nat Commun 10:3627.

8. Fan SH, Liberini E, Gotz F. 2021. *Staphylococcus aureus* Genomes Harbor Only MpsAB-Like Bicarbonate Transporter but Not Carbonic Anhydrase as Dissolved Inorganic Carbon Supply System. Microbiol Spectr 9:e0097021.

9. Nishida H, Beppu T, Ueda K. 2009. *Symbiobacterium* lost carbonic anhydrase in the course of evolution. J Mol Evol 68:90–6.

10. Perez-Etayo L, de Miguel MJ, Conde-Alvarez R, Munoz PM, Khames M, Iriarte M, Moriyon I, Zuniga-Ripa A. 2018. The CO(2)-dependence of *Brucella ovis* and *Brucella abortus* biovars is caused by defective carbonic anhydrases. Vet Res 49:85.

11. Varesio LM, Willett JW, Fiebig A, Crosson S. 2019. A Carbonic Anhydrase Pseudogene Sensitizes Select *Brucella* Lineages to Low CO(2) Tension. J Bacteriol 201.

12. Garcia Lobo JM, Ortiz Y, Gonzalez-Riancho C, Seoane A, Arellano-Reynoso B, Sangari FJ. 2019. Polymorphisms in *Brucella* Carbonic Anhydrase II Mediate CO(2) Dependence and Fitness in vivo. Front Microbiol 10:2751.

13. Jiang M, Chen M, Guo ZF, Guo Z. 2010. A bicarbonate cofactor modulates 1,4-dihydroxy-2-naphthoyl-coenzyme a synthase in menaquinone biosynthesis of *Escherichia coli*. J Biol Chem 285:30159–69.

14. Aguilera J, Van Dijken JP, De Winde JH, Pronk JT. 2005. Carbonic anhydrase (Nce103p): an essential biosynthetic enzyme for growth of *Saccharomyces cerevisiae* at atmospheric carbon dioxide pressure. Biochem J 391:311–6.

15. Wille W, Eisenstadt E, Willecke K. 1975. Inhibition of de novo fatty acid synthesis by the antibiotic cerulenin in *Bacillus subtilis*: effects on citrate-Mg2+ transport and synthesis of macromolecules. Antimicrob Agents Chemother 8:231–7.

16. Nickels JD, Chatterjee S, Stanley CB, Qian S, Cheng X, Myles DAA, Standaert RF, Elkins JG, Katsaras J. 2017. The in vivo structure of biological membranes and evidence for lipid domains. PLoS Biol 15:e2002214.

17. Elfmann C, Dumann V, van den Berg T, Stulke J. 2025. A new framework for SubtiWiki, the database for the model organism *Bacillus subtilis*. Nucleic Acids Res 53:D864–D870.

18. Schau M, Eldakak A, Hulett FM. 2004. Terminal oxidases are essential to bypass the requirement for ResD for full Pho induction in *Bacillus subtilis*. J Bacteriol 186:8424–32.

19. Valley G, Rettger Leo F. 1927. The influence of carbon dioxide on bacteria. Journal of Bacteriology 14:101–137.

20. Gladstone GP, Fildes P, Richardson GM. 1935. Carbon Dioxide as an Essential Factor in the Growth of Bacteria. Br J Exp Pathol 16:335–48.

21. Bennett BD, Kimball EH, Gao M, Osterhout R, Van Dien SJ, Rabinowitz JD. 2009. Absolute metabolite concentrations and implied enzyme active site occupancy in *Escherichia coli*. Nat Chem Biol 5:593–9.

22. Libor SM, Sundaram TK, Scrutton MC. 1978. Pyruvate carboxylase from a thermophilic *Bacillus*. Studies on the specificity of activation by acyl derivatives of coenzyme A and on the properties of catalysis in the absence of activator. Biochem J 169:543–58.

23. Brugarolas P, Duguid EM, Zhang W, Poor CB, He C. 2011. Structural and biochemical characterization of N5-carboxyaminoimidazole ribonucleotide synthetase and N5-carboxyaminoimidazole ribonucleotide mutase from *Staphylococcus aureus*. Acta Crystallogr D Biol Crystallogr 67:707–15.

24. Fan SH, Matsuo M, Huang L, Tribelli PM, Gotz F. 2021. The MpsAB Bicarbonate Transporter Is Superior to Carbonic Anhydrase in Biofilm-Forming Bacteria with Limited CO(2) Diffusion. Microbiol Spectr 9:e0030521.

25. Oppenheimer-Shaanan Y, Sibony-Nevo O, Bloom-Ackermann Z, Suissa R, Steinberg N, Kartvelishvily E, Brumfeld V, Kolodkin-Gal I. 2016. Spatio-temporal assembly of functional mineral scaffolds within microbial biofilms. NPJ Biofilms Microbiomes 2:15031.

26. Kim JK, Lee C, Lim SW, Adhikari A, Andring JT, McKenna R, Ghim CM, Kim CU. 2020. Elucidating the role of metal ions in carbonic anhydrase catalysis. Nat Commun 11:4557.

27. Bhandary D, de Visser SP, Mukherjee G. 2025. Implications of non-native metal substitution in carbonic anhydrase - engineered enzymes and models. Chem Commun (Camb) 61:612–626.

28. Hartig E, Jahn D. 2012. Regulation of the anaerobic metabolism in *Bacillus subtilis*. Adv Microb Physiol 61:195–216.

29. Michna RH, Commichau FM, Todter D, Zschiedrich CP, Stulke J. 2014. SubtiWiki-a database for the model organism *Bacillus subtilis* that links pathway, interaction and expression information. Nucleic Acids Res 42:D692–8.

30. Kreuzer-Martin HW, Ehleringer JR, Hegg EL. 2005. Oxygen isotopes indicate most intracellular water in log-phase *Escherichia coli* is derived from metabolism. Proceedings of the National Academy of Sciences 102:17337–17341.

31. Jacobson TA, Kler JS, Hernke MT, Braun RK, Meyer KC, Funk WE. 2019. Direct human health risks of increased atmospheric carbon dioxide. Nature Sustainability 2:691–701.

32. Koo BM, Kritikos G, Farelli JD, Todor H, Tong K, Kimsey H, Wapinski I, Galardini M, Cabal A, Peters JM, Hachmann AB, Rudner DZ, Allen KN, Typas A, Gross CA. 2017. Construction and Analysis of Two Genome-Scale Deletion Libraries for *Bacillus subtilis*. Cell Syst 4:291–305 e7.

33. Soding J, Biegert A, Lupas AN. 2005. The HHpred interactive server for protein homology detection and structure prediction. Nucleic Acids Res 33:W244–8.

34. Madeira F, Madhusoodanan N, Lee J, Eusebi A, Niewielska A, Tivey ARN, Lopez R, Butcher S. 2024. The EMBL-EBI Job Dispatcher sequence analysis tools framework in 2024. Nucleic Acids Res 52:W521–W525.

35. Meng EC, Goddard TD, Pettersen EF, Couch GS, Pearson ZJ, Morris JH, Ferrin TE. 2023. UCSF ChimeraX: Tools for structure building and analysis. Protein Sci 32:e4792.

36. Jumper J, Evans R, Pritzel A, Green T, Figurnov M, Ronneberger O, Tunyasuvunakool K, Bates R, Zidek A, Potapenko A, Bridgland A, Meyer C, Kohl SAA, Ballard AJ, Cowie A, Romera-Paredes B, Nikolov S, Jain R, Adler J, Back T, Petersen S, Reiman D, Clancy E, Zielinski M, Steinegger M, Pacholska M, Berghammer T, Bodenstein S, Silver D, Vinyals O, Senior AW, Kavukcuoglu K, Kohli P, Hassabis D. 2021. Highly accurate protein structure prediction with AlphaFold. Nature 596:583–589.

37. Schindelin J, Arganda-Carreras I, Frise E, Kaynig V, Longair M, Pietzsch T, Preibisch S, Rueden C, Saalfeld S, Schmid B, Tinevez JY, White DJ, Hartenstein V, Eliceiri K, Tomancak P, Cardona A. 2012. Fiji: an open-source platform for biological-image analysis. Nat Methods 9:676–82.

